# The ant fungus garden acts as an external digestive system

**DOI:** 10.1101/2020.11.18.389361

**Authors:** Andrés Mauricio Caraballo-Rodríguez, Sara P. Puckett, Kathleen E. Kyle, Daniel Petras, Ricardo da Silva, Louis-Félix Nothias, Madeleine Ernst, Justin J.J. van der Hooft, Anupriya Tripathi, Mingxun Wang, Marcy J. Balunas, Jonathan L. Klassen, Pieter C. Dorrestein

## Abstract

Most animals digest their food within their own bodies, but some do not. Many species of ants grow fungus gardens that pre-digest food as an essential step of the ants’ nutrient uptake. To better understand this digestion process, we generated a 3D molecular map of an *Atta texana* fungus garden, revealing chemical modifications mediated by the fungus garden as plant material passes through.

## Main

Many ant species access plant-derived nutrients with the help of fungal symbionts.^1^ For example, leaf-cutter ants grow a specific cultivar fungus in specialized underground structures called “fungus gardens” as their main food source. This cultivar fungus breaks down forage material such as leaves provided by the ants to obtain the necessary nutrients for its own growth.^2^ In turn, the ants eat the fungus’ specialized hyphal tips, known as gongylidia, which contain nutrients that are metabolically available to the ants.^3^ Fungal enzymes present in the garden transform plant metabolites such as polysaccharides and phenolic compounds.^4–6^ Fungus gardens from leaf-cutter ants have been described as bioreactors due to their capacity to process plant constituents that provide small carbohydrates.^6^ Primary metabolites have been measured in the fungus gardens, and correlated to the differential distribution of fungal metabolic enzymes.^2,7^ Nonetheless, maps of metabolic diversity in ant fungus gardens have remained unavailable due to the lack of computational workflows that go beyond the analysis of a few selected metabolites, which did not exist until recently.

Here, we highlight chemical transformations in a laboratory maintained *Atta texana* fungus garden using molecular networking,^8–10^ 3D cartography,^11^ and meta mass-shift analysis^12^. The use of non-targeted metabolomics, via liquid chromatography – tandem mass spectrometry (LC-MS/MS),^10,13–15^ enabled us to identify molecular families and metabolite features (**Supplementary Fig. S1-S2**) that chemically differentiate plant materials that are sequentially consumed by the fungus as they pass through the garden (**Supplementary Fig. S3-S4**). These molecular families were annotated using various annotation tools combined using the MolNetEnhancer^10^,^16^ workflow resulting in the annotation of plant and fungus related chemical compound classes. Furthermore, we identified the types of chemical transformations that are carried out based on the differential abundance of compounds that occur among the sampled layers of the fungus garden (**Online methods**). The observed transformations provide insight into the chemistry and the modification of molecules (representing potential chemical transformations or differential degradation) inside an ant fungus garden.

After plant materials are incorporated into the fungus garden by the ants, they are further processed by the fungus, and following digestion any recalcitrant plant biomass becomes trash that the ants remove from the garden (**Fig. 1**). Molecules from the plant material, such as saccharide-decorated flavonoids and phenolic compounds, decreased in relative abundance when moving from the top to the bottom of the fungus garden, in contrast to other compounds that increased in relative abundance across these layers (**Supplementary Fig. S3**), either due to chemical modifications or preferential degradation of the less abundant compound (**Supplementary Fig. S5-S14**). Fungus garden and trash samples were enriched with phytosphingosines (**Supplementary Fig. S9**), whereas features associated with the trash material were enriched in steroids (**Fig. 1, Supplementary Fig. S10**), such as the fungal metabolite ergosterol peroxide,^17,18^ and oxylipins (**Supplementary Fig. S11-S12**). Gradients of other plant-derived metabolite abundances, such as triterpenoid derivatives, were observed as we moved from the top to the bottom of the fungus garden, leading to high abundances at the bottom of the fungus garden and in the trash (**Supplementary Fig. S13-S14**). These gradients parallel the metabolic transformations of food components in the digestive tract of animals, such as those involving the metabolism of flavonoids, steroids (molecules with steroidal cores), and fatty acids.^19–21^ In a similar way to how food changes during its transit through the digestive tract of animals and the residual material is discarded, plant material is transformed in the fungus garden and, finally, the residual material is removed from the colony. Thus, the initial food material is chemically distinct when compared to the trash material removed by the ants (**Supplementary Fig. S4**).

**Fig. 1.**
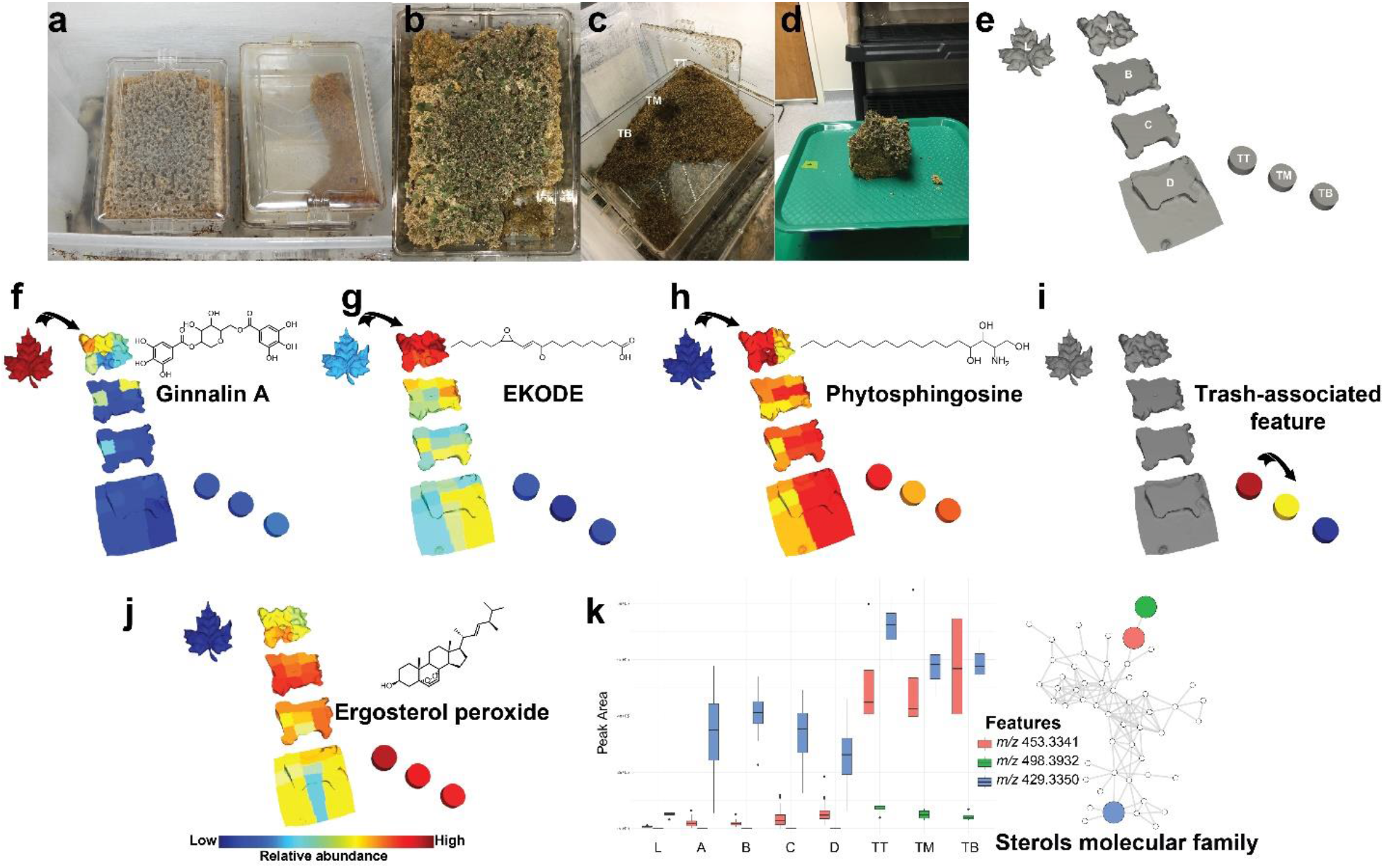
Spatial distribution of molecular signatures from *A. texana* fungus garden. **a-d.** Deconstruction of the *Atta texana* JKH000189 fungus garden: **a.** Plastic containers with an *Atta texana* fungus garden (left) and waste material removed by ants (right). Ants have free access to both chambers; **b.** This picture shows the location of a removed 10×10×10cm portion of the fungus garden (lower right corner). The green fragments at the top of the fungus garden are freshly incorporated maple leaves; **c.** Chamber containing the trash material removed from the fungus garden by the ants. Three sampling locations from the trash are highlighted as “TB”, “TM”, and “TT”; **d.** Removed 10×10×10cm fungus garden portion for consequent sample preparation for LC-MS/MS; **e.** 3D representation of deconstructed fungus garden portion, as shown in (**d**). On the left side, a representation of the maple leaves (L) placed in the outer colony box that ants cut and incorporate into the top of the fungus garden; layers of the fungus garden from top to bottom (A, B, C, and D) and at the right side of the figure, the representation of the three sample locations from the trash chamber, from top to bottom (TT, TM and TB); **f-i.** Spatial distribution of ginnalin A, detected as *m/z* 469.0971 (**f**), (E)-9-oxo-11-(3-pentyloxiran-2-yl)undec-10-enoic acid (trans-EKODE-(E)-lb) detected as *m/z* 311.221 (**g**), phytosphingosine detected as *m/z* 318.2995 (**h**), unknown feature associated to the trash material detected as *m/z* 474.3783 (**i**); **j-k.** The abundance of features belonging to the molecular family of sterols were also detected at high intensity in the trash material, suggesting that these compounds accumulate in the trash material: **j.** Ergosterol peroxide detected as *m/z* 429.3350; **k.** Abundance of features associated with trash material belonging to the sterols molecular family (ergosterol peroxide, feature *m/z* 498.3932 and feature *m/z* 453.3341). The boxes represent the 25%, 50%, and 75% quantile and the whiskers extend ±1.5 times the interquartile range. The annotation of ergosterol peroxide from GNPS libraries (cosine score = 0.71) was confirmed using a reference standard (**Supplementary Table S1**), a level 1 match according to the 2007 metabolomics initiative,^32^ while the *m/z* 498.3932 is consistent with a molecular formula of C_32_H_52_NO_3_ (error 1.9 ppm) and belongs to the same molecular family – a level 3 match.^32^ A detailed description of the sample preparation can be found in the **Online methods.** See **online methods** for more details and a molecular cartography of this deconstructed *Atta texana* fungus garden visualized in ‘ili^11^ is shown in **Supplementary Movie S1 [https://youtu.be/_ikhKelfrY8]**.

Digestive processes generate modified products whose precursors are consumed. To provide an overview of putative metabolic transformations occurring in ants’ fungus gardens, we combined mass shift analysis^12^ and discovered relative metabolite abundances for pairs from each section of the fungus garden by calculating a proportionality score (**Online Methods**). By considering the proportion between the relative abundances of two chemically related molecules (i.e. connected nodes in a network), their mass shifts and the modifications that these imply (e.g. a 15.996 Da shift indicates a gain or loss of oxygen, 2.015 Da an oxidation or reduction through the loss or addition of H2), and their distribution between two locations (leaves, layers of fungus garden and layers of trash material) we can discover related molecules that have the largest variance in abundance between the layers (**Online Methods**, **Fig. 1–3; Supplementary Fig. S15-S17)**. It should be noted that this approach cannot differentiate between different types of changes in the absolute abundance of each molecule, e.g., chemical transformation leading to the accumulation of a molecule or the complete degradation of a molecule leading to its decreased abundance. However, by considering the chemical similarities and relative abundances between each molecular pair that are identified by molecular networking we imply relationships between these molecules that are consistent with each molecule belonging to the same structural class and their abundance changes across samples. We interpret the abundance changes across layers to be largely driven by anabolic or catabolic pathways, potentially linked to enzymatically mediated transformations. Absolute molecular concentrations might also be altered by additions from the external environment, but the closed nature of the laboratory maintained ant fungus gardens means that such additions essentially only occur directionally via the leaves when they are incorporated into the fungus garden by the ants.

**Fig. 2|.**
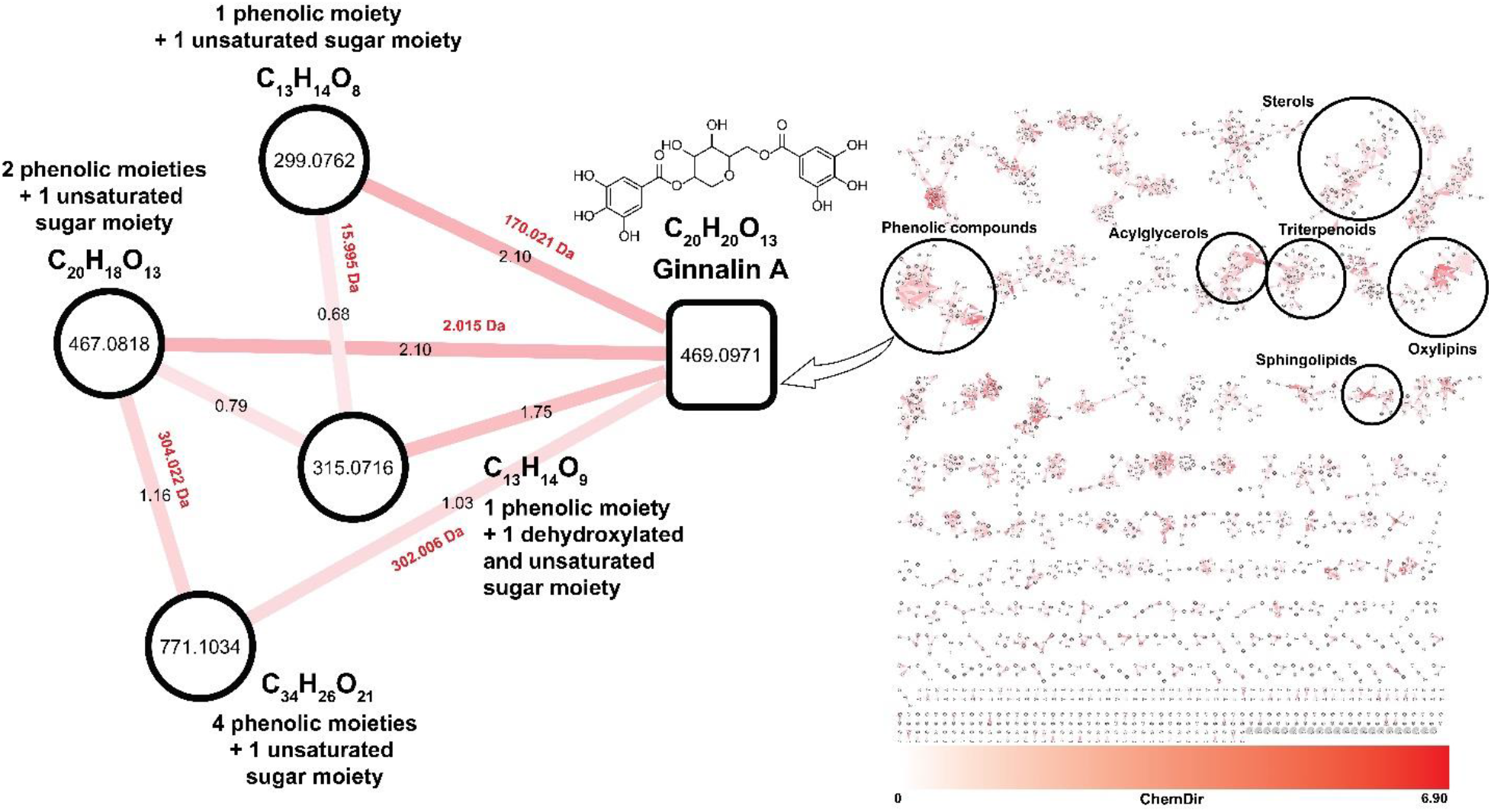
Potential transformations highlighted by the proportional scoring of untargeted metabolomics data from *Atta texana* fungus garden. Chemical features are highlighted from a molecular network based on their high proportionality score calculated throughout the sample types from leaves, fungus garden layers and trash layers (red edge with label indicating the proportionality score). The feature corresponding to *m/z* 469.0971 annotated as ginnalin A, a bioactive phenolic compound,^25^ was identified as a potential partner in several chemical transformations. Based on the information regarding the mass shifts (red labels, in Daltons) of the connected nodes, and the suggested molecular formula, putative structures can be suggested based on their fragmentation patterns. Together, with the spatial distribution (**Supplementary Figures S8 and S16**), it can be suggested the involvement of this molecular family in chemical transformations occurring in the fungus garden. Ginnalin A was detected in the leaves as well as in the fungus garden while the features of *m/z* 299.0762 and *m/z* 467.0818 were only detected in the plant material. This suggest that ginnalin A and related features, *m/z* 315.0717 and *m/z* 771.1034 might be either transformed from *m/z* 467.0818 and *m/z* 299.0762, which reach undetectable levels in the fungus garden, or are more recalcitrant to degradation. Similarities in the fragmentation spectra (MS/MS) of these compounds enabled us to gain insight into their potential structural differences. All fragmentation spectra for the connected nodes in the molecular network shown in the figure shared the *m/z* 153.02 base peak corresponding to the phenolic moiety (gallic acid), indicating an unsaturation located in the sugar moiety. The identification of ginnalin A based on spectral similarity to GNPS libraries (cosine score = 0.95) corresponds to annotation at level 2 according to the 2007 metabolomics standards initiative,^26^ while the matches for the related molecules shown are at the molecular family level, a level 3 annotation.^26^ Although double bonds of the saccharide moiety can be suggested based on the molecular formula, it is not possible to define their location or the stereochemistry thereof without additional information. Other examples of potential chemical transformations in *Atta texana* fungus gardens described using this proportionality approach can be found in **Supplementary Fig. S15-S17)**.

**Fig. 3|.**
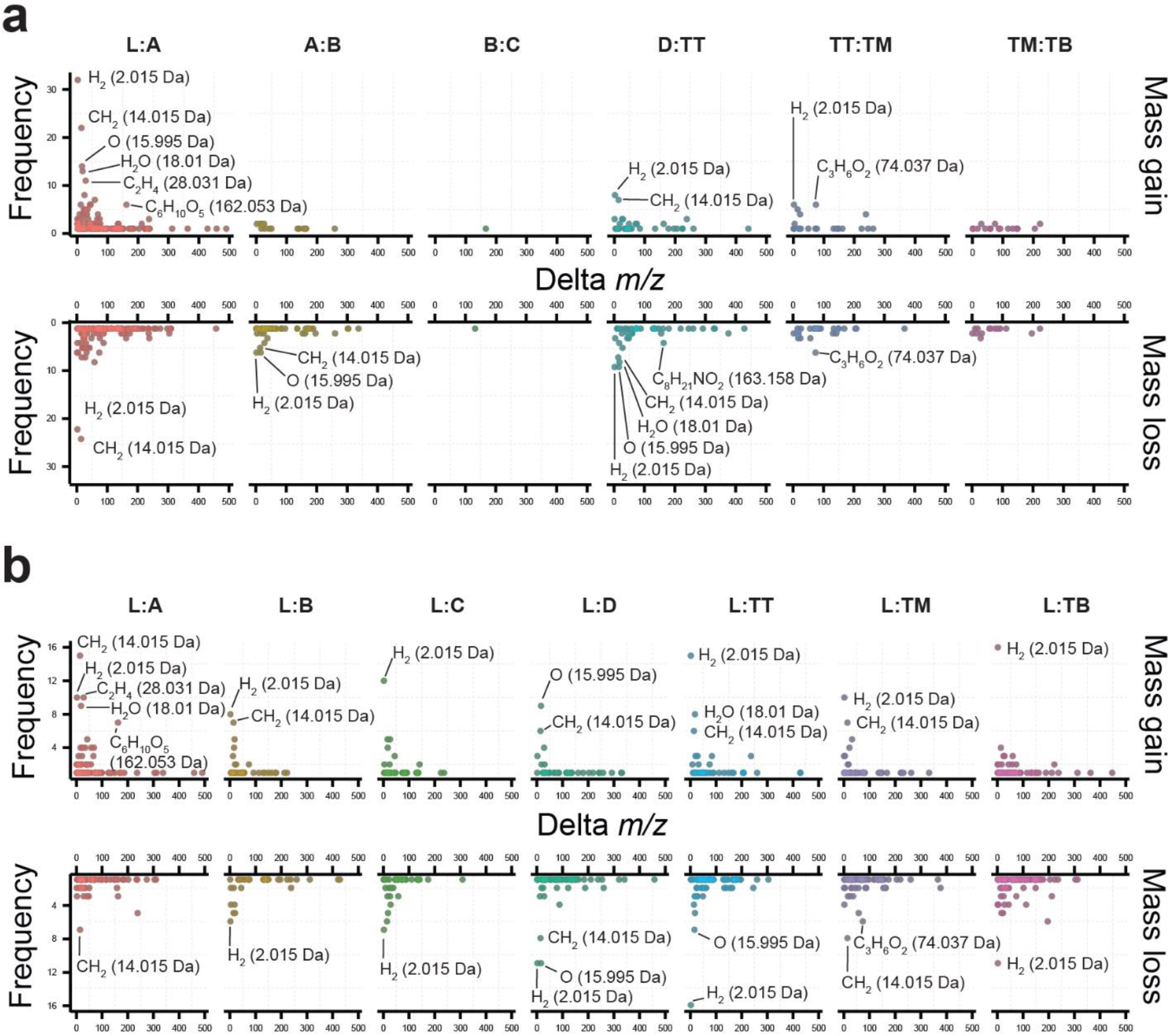
Frequency of delta masses observed in metabolomics data from a deconstructed *Atta texana* fungus garden. Proportionality metrics were calculated between compounds found in different layers of the fungus garden, as well as between plant, fungus garden and trash material: **a.** Frequency of delta masses calculated between samples corresponding to leaves (L), layers of fungus garden from top to bottom (A, B, C and D), and trash material (samples collected from the chamber that ants use to deposit the trash material. From the top, TT; middle, TM and bottom, TB). **b.** Frequency of delta masses between compounds found in leaves (L) and each of the fungus garden and trash layers. Mass shifts between network pairs with proportionality scores > 1 were retrieved as indicative of chemical transformations prevalent in specific locations. Frequencies of annotated mass shifts observed from network pairs with high proportionality scores (proportionality values > 1, See **Supplementary Table S4**) indicated that the regions where most transformations occur are between leaves and the top layer of fungus garden (Layer A), between the bottom of fungus garden (Layer D) and the trash, as well as between the trash layers (TT, TM and TB). None of the observed mass shifts resulted in a high score when proportionality was calculated between fungus layers C and D (proportionality C:D). The figure in panel **b** shows the most frequent mass shifts observed from network pairs with highest transformation rates from proportionality calculated between leaves (L) and each of the sample groups, fungus garden and trash layers. This indicates that most of these transformations are still observed throughout the fungus garden but also that some of them are specific to the trash, such as C_8_H_21_NO_2_ (163.158 Da) (as observed in panel **a**).

In this study, we introduce the proportionality concept using a modification to Meta-Mass Shift analysis,^12^ by considering also the abundances of the detected molecules, to quickly highlight features (metabolites) potentially involved in chemical transformations. Proportionality scores highlighted such changing pairs of nodes occurring in various locations in the *A. texana* fungus garden, and the associated mass shifts provided evidence of the types of modifications occurring in the sampled locations (**Fig. 2**). Proportionality scoring is a logarithmic expression, and we selected an absolute value of 1 as cutoff to prioritize potential transformations because a score closes to zero will indicate a low ratio of abundances between two chemically related molecules in two sample locations, suggesting a low change of intensities between the two mass spectrometry features of the molecular network pair (see the definition and calculation of the proportionality scores in **Online Methods**). Differences representing a gain or loss of H2 (2.015 Da) were the predominant type of chemical transformation observed throughout the entire data set, being one of the most frequent mass shifts with a proportionality score > 1 among leaves, the fungus garden and trash layers (**Fig. 3**). This common modification was observed in molecular families such as phenolic compounds (**Fig. 2**) and phytosphingosines (**Supplementary Fig. S9**).^22^ Mass shifts corresponding to CH2 (14.015 Da) and C2H4 (28.031 Da) were other common changes observed in the top layer of the fungus garden, as well as between the bottom layer of the fungus garden and the trash (**Fig. 3**). Oxidation or dehydroxylation combined with reduction processes result in the gain or loss of oxygen that can be detected and a mass difference of 15.995 Da. The transformations corresponding to these differences were more frequently observed at the top layer of the fungus garden during the breakdown of flavonoids and phenolic compounds (**Fig. 3, Supplementary Fig. S5-S8**).

Chemical transformations consistent with addition or removal of sugar moieties in the *A. texana* fungus garden were also highlighted by proportionality scores > 1. These transformations, corresponding to mass differences of 162.053 Da (C6H10O5), were associated with plant material and the top layers of the fungus garden (**Fig. 3**), and corresponded to transformations involving chemical substructures that were present in plant metabolites such as flavonoids and acylglycerols (**Supplementary Fig. S6, S17**). The mass difference of 162.053 Da, consistent with the gain or loss of a sugar moiety, was also present in a molecular family of acylglycerols and occurred in the top layer of the fungus garden (**Supplementary Fig. S17**).

The existence of chemical gradients in the fungus garden resembles a digestion process, with substrates being modified to facilitate their consumption and generating residues that need to be removed or discarded, as shown here by plant constituents passing through an ant-fungus garden ecosystem. The metabolic transformations observed here are consistent with the modification of lipids by fungus gardens, as well as plant volatile compounds by fungus-garden-associated bacteria, as recently reported from leaf-cutter ant fungus gardens,^23,24^ adding further support to the model describing fungus gardens as external digestive systems for ants. The chemical modifications and the types of (bio)transformations observed in our study might not vary based on changes in the available plant material, although environmental factors such as temperature or humidity, and the composition of microbiome that are associated with ants and their fungus gardens,^23^ will likely influence these modifications.

In summary, the 3D cartographic analyses performed in this study provide an overview of chemical changes occurring in a fungus garden. Our results demonstrate that chemical transformations of the plant components are associated with certain regions of the fungus garden and show that the degree of modifications are more extensive than previously described.^24^,^25^ The results further support a model where fungus gardens serve a similar function to the mammalian digestive tract, where its function is to gradually metabolize food molecules from top to bottom akin to the gastrointestinal tract from the mouth to anus. In other words, the plant material is digested starting when leaves enter the fungus garden, and continues all the way through the bottom layer of the fungus garden. Finally, there are molecules that are removed from the system into the fungus garden trash. This is reminiscent of a food to digestive tract to feces scheme.^21^ How the food molecules move down the ant’s external digestive tract is not yet known, but the ants “clean up” after themselves via removal of unwanted fungal garden parts to the trash. The methodologies that we used provide a complementary overview of metabolic processes occurring in a laboratory maintained *A. texana* fungus garden and we expect the approach can be leveraged to unravel similar processes in natural environments to compare between natural and lab-maintained ecosystems.

## Online content

Methods, additional references, Nature Research reporting summaries, source data, statements of data availability and associated accession codes are available on line.

## Acknowledgements

AMCR and PCD were supported by the National Sciences Foundation grant IOS-1656481. KEK, SPP, JLK and MJB were supported by NSF grant IOS-1656475. DP was supported by the Deutsche Forschungsgemeinschaft (DFG) with grant PE 2600/1. RRdS was supported by the São Paulo Research Foundation (Awards FAPESP 2017/18922–2 and 2019/05026–4). PCD was supported by the Gordon and Betty Moore Foundation through Grant GBMF7622, the U.S. National Institutes of Health for the Center (P41 GM103484, R03 CA211211, R01 GM107550). LFN was supported by the U.S. National Institutes of Health (R01 GM107550). JJJvdH was supported by an ASDI eScience grant, ASDI.2017.030, from the Netherlands eScience Center — NLeSC. We thank Alan K. Jarmusch for his valuable comments in earlier versions of the manuscript.

