## Supplementary material for "The ant fungus garden acts as an external digestive system": Online Methods and Supplementary Information

**Authors:** Andrés Mauricio Caraballo-Rodríguez[1], Sara P. Puckett[2], Kathleen E. Kyle[3], Daniel Petras[1,4], Ricardo da Silva[5], Louis-Félix Nothias[1], Madeleine Ernst[6], Justin J.J. van der Hooft[7], Anupriya Tripathi[1,8,9], Mingxun Wang[1,10], Marcy J. Balunas[2], Jonathan L. Klassen[3]**, Pieter C. Dorrestein[1]*

**Affiliations**

[1] Collaborative Mass Spectrometry Innovation Center, Skaggs School of Pharmacy and Pharmaceutical Sciences, University of California San Diego, La Jolla, CA.

[2] Division of Medicinal Chemistry, Department of Pharmaceutical Sciences, University of Connecticut, Storrs, CT 06269, USA.

[3] Department of Molecular and Cell Biology, University of Connecticut, Storrs, CT, USA.

[4] Scripps Institution of Oceanography, University of California San Diego, La Jolla, CA

[5] School of Pharmaceutical Sciences of Ribeirão Preto, University of São Paulo, Ribeirão Preto, SP, Brazil.

[6] Section for Clinical Mass Spectrometry, Danish Center for Neonatal Screening, Department of Congenital Disorders, Statens Serum Institut, Copenhagen, Denmark

[7] Bioinformatics Group, Wageningen University, 6708 PB, Wageningen, the Netherlands.

[8] Division of Biological Sciences, University of California—San Diego, San Diego, California, USA

[9] Department of Pediatrics, University of California—San Diego, San Diego, California, USA

[10] Ometa Labs LLC

* to whom correspondence should be addressed for the mass spectrometry and data analysis:.** to whom correspondence should be addressed regarding the ant-fungus system:.

### Supplementary Figures

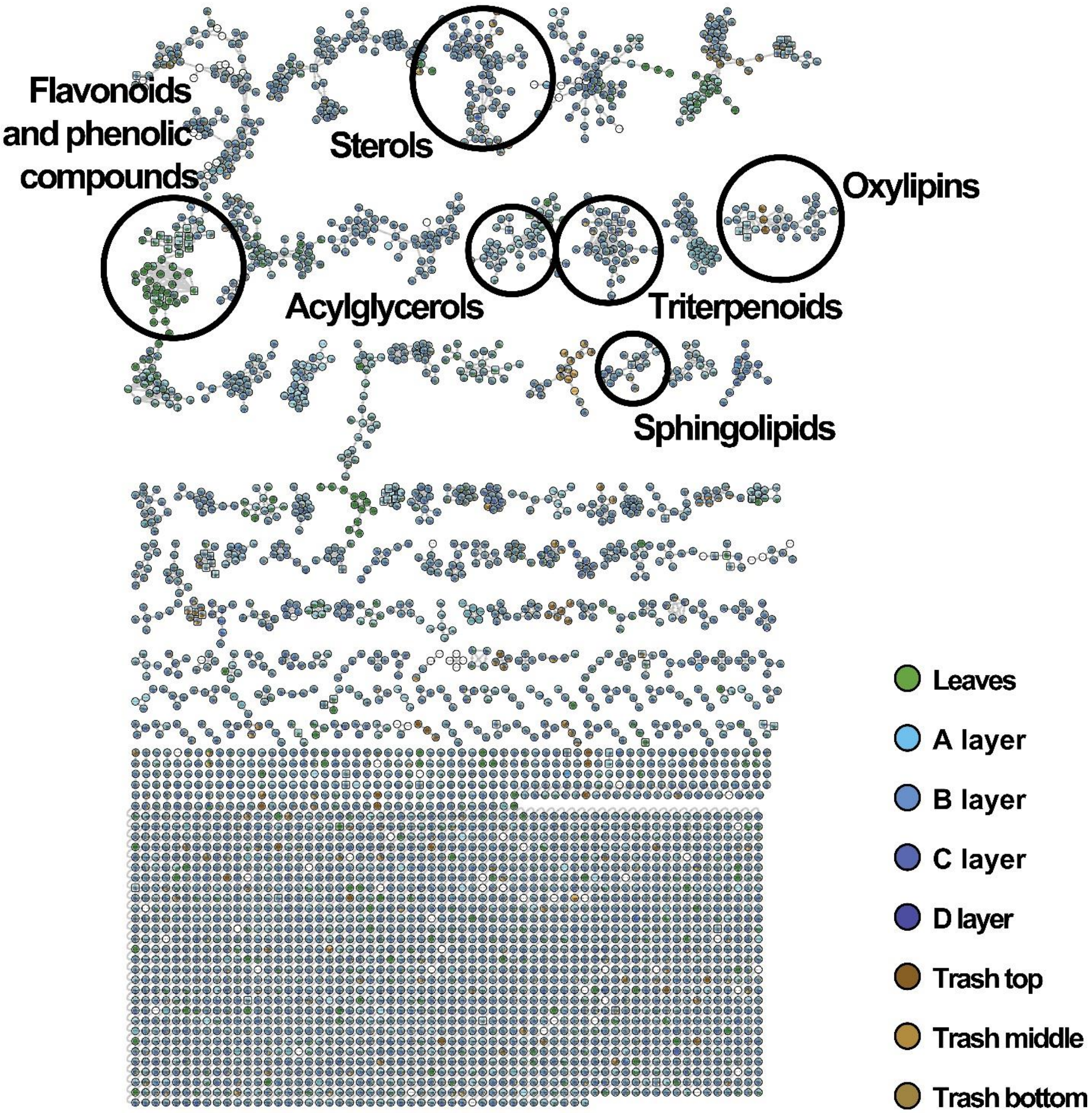


**Supplementary Fig. S1|Molecular network from deconstructed *Atta texana* fungus garden.** Molecular network highlighting molecular families containing putatively identified compounds from GNPS spectral libraries. Connected nodes correspond to structurally related molecules based on their spectral similarity. Color codes as described in the figure. The molecular networking job can be accessed via the following link: [https://gnps.ucsd.edu/ProteoSAFe/status.jsp?task=5df1dc83e075478ba69d1bb41bf9499a].

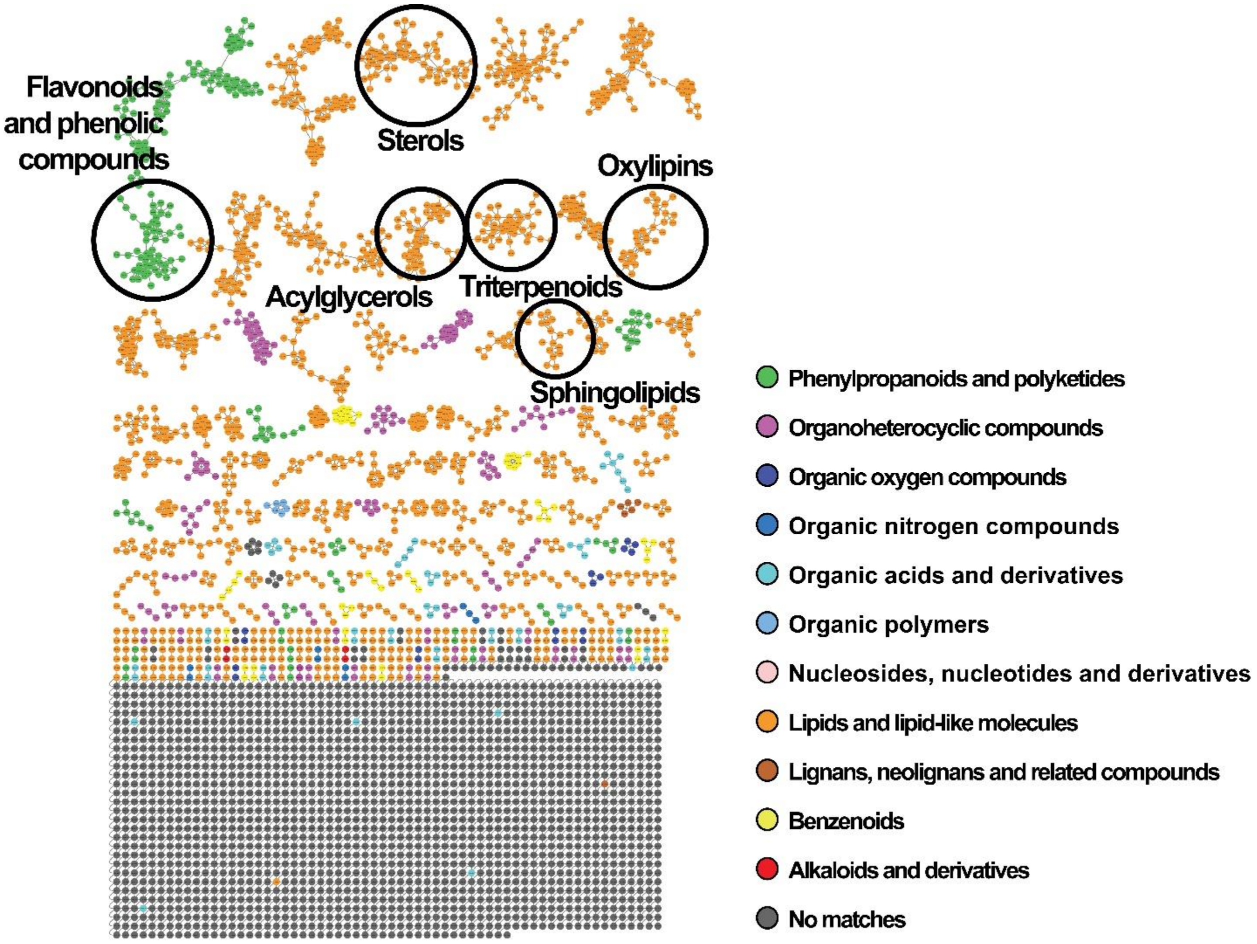


**Supplementary Fig. S2|Enhanced annotation of molecular network from deconstructed *Atta texana* fungus garden.** Molecular network including expanded annotations for detected features via *in silico* tools using the MolNetEnhancer[1] workflow available in the GNPS platform. Color codes as described in the figure represent the most predominant putative annotation of the detected features per molecular family at the superclass chemical taxonomy level according to Feunang *et al*., 2016.[2] Additionally, the molecular families containing putatively identified compounds from GNPS spectral libraries are highlighted and labeled. The enhanced molecular networking job can be accessed via the following link: https://gnps.ucsd.edu/ProteoSAFe/status.jsp?task=50dfaf589c8140bd85a8ca199db59de3

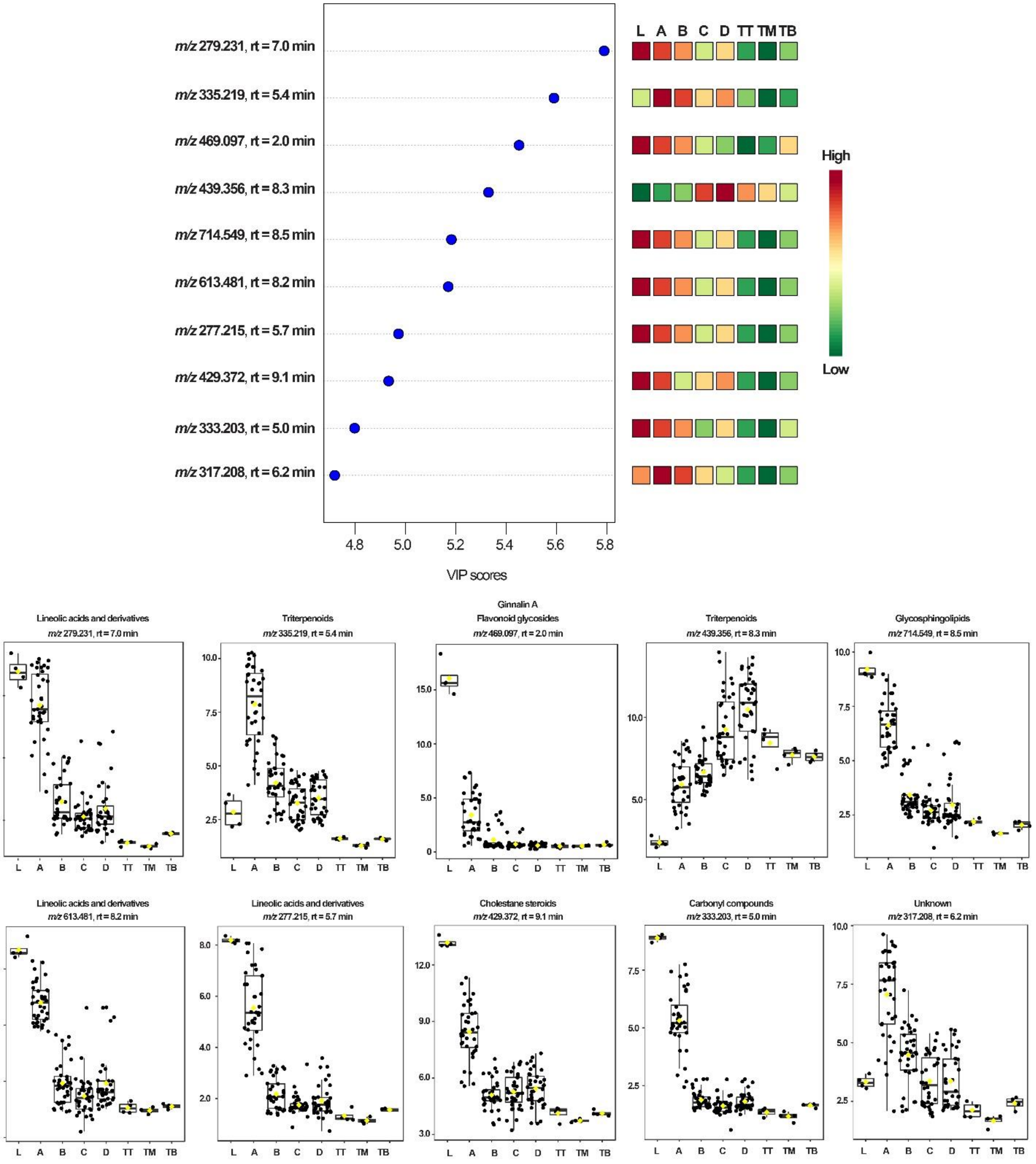

m/z 279.231, rt = 7.0 min
m/z 335.219, rt = 5.4 min
m/z 469.097, rt = 2.0 min
m/z 439.356, rt = 8.3 min
m/z 714.549, rt = 8.5 min
m/z 613.481, rt = 8.2 min
m/z 277.215, rt = 5.7 min
m/z 429.372, rt = 9.1 min
m/z 333.203, rt = 5.0 min
m/z 317.208, rt = 6.2 min
4.8
5.0
5.2
5.4
5.6
5.8
VIP scores
L A B C D TT TM TB
High
Low
Lineolic acids and derivatives
Triterpenoids
Ginnalin A
Flavonoid glycosides
Triterpenoids
Glycosphingolipids
Lineolic acids and derivatives
Lineolic acids and derivatives
Cholestane steroids
Carbonyl compounds
Unknown

**Supplementary Fig. S3|Chemical features with high contribution in differentiation of samples from *Atta texana* fungus garden.** Separation of layers of the fungus garden is clearly observed between the top layer (A layer) against the other fungus layers (B, C and D layers). From the top 10 features with high loadings score (VIP score) from PLS-DA are shown with annotation at the subclass level based on candidates from NAP[3] or CSI:FingerID[4] and Classyfire[2] chemical classification approach were no identification provided by GNPS libraries, otherwise the library match is specified at the top. The whisker plots for the detected features correspond to normalized values using internal standard (sulfamethazine) as reference feature (y-axis). Samples, as shown in x-axis correspond to the following labels: maple leaves (L); layers of the fungus garden from top to bottom (A, B, C, and D) and trash from top to bottom (TT, TM and TB). Statistical analysis performed using MetaboAnalyst platform.[5,6]

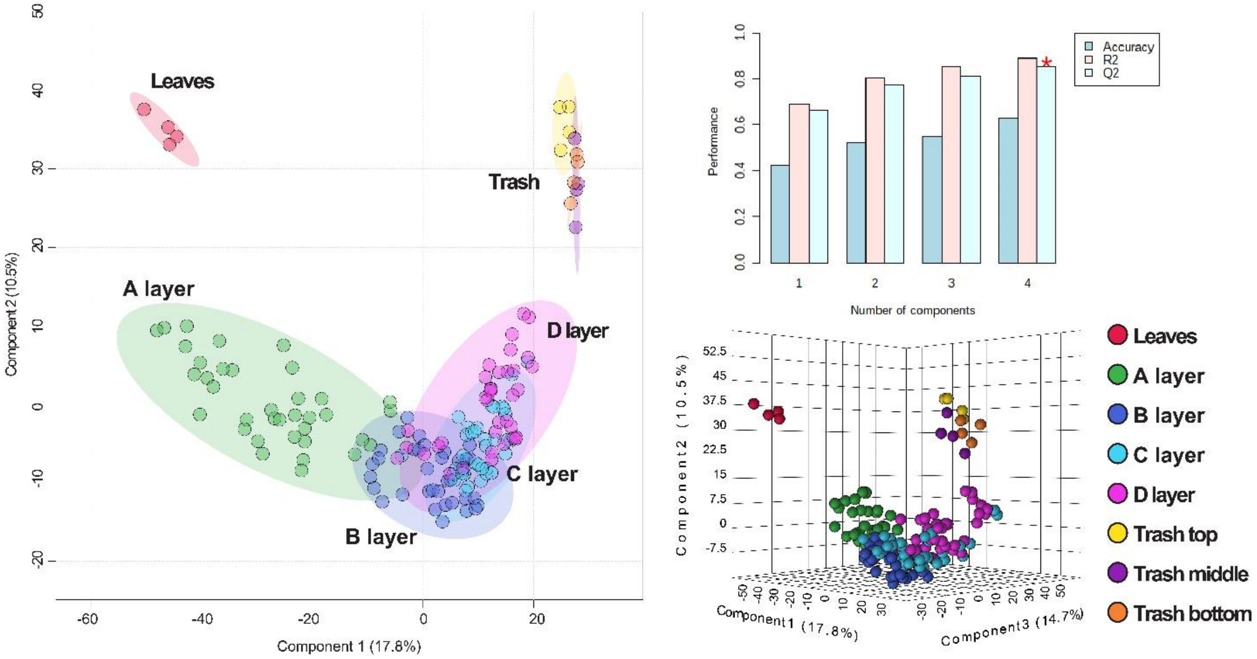


**Supplementary Fig. S4|Chemical difference of *Atta texana* fungus garden samples via PLS-DA.** PLS-DA plots obtained from statistical analysis using quantification tables containing detected features from *Atta texana* fungus garden. Evident separation was obtained for leaves and trash material when compared against fungus garden layers. Statistical analysis performed using MetaboAnalyst platform.[5,6] PLS-DA outputs (coef., loadings, scores and vip scores) used to generate the corresponding plots are provided as **Supplementary Table S4-S7**.

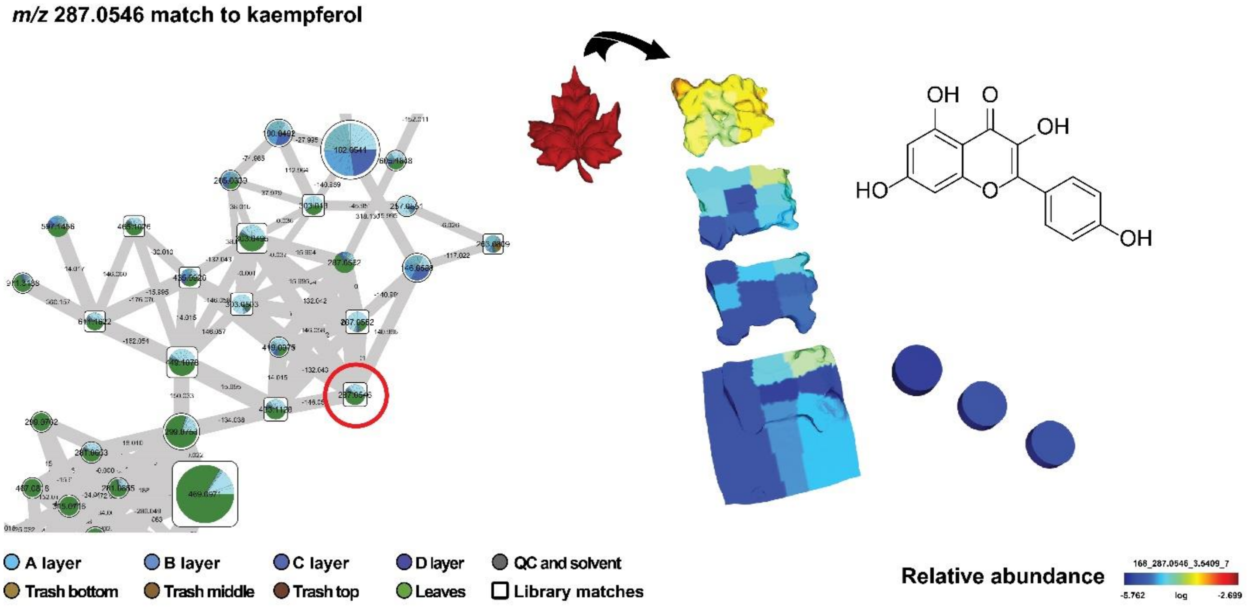


**Supplementary Fig. S5|Molecular cartography of kaempferol detected in *A. texana* fungus garden.** In this figure, the detected feature corresponding to kaempferol (m/z 287.0546) is highly abundant in the plant material and in the top layer of the fungus garden when compared to the remaining layers of fungus garden and trash material. Connected nodes correspond to structurally related molecules based on their spectral similarity and due to their distribution (mainly fungus garden), they might be products of biotransformation processes occurring in the fungus garden. The annotation of kaempferol from GNPS libraries (cosine 0.93) was confirmed by using a reference standard (**Supplementary Table S1**), a level 1 annotation according to the 2007 metabolomics initiative.[7]

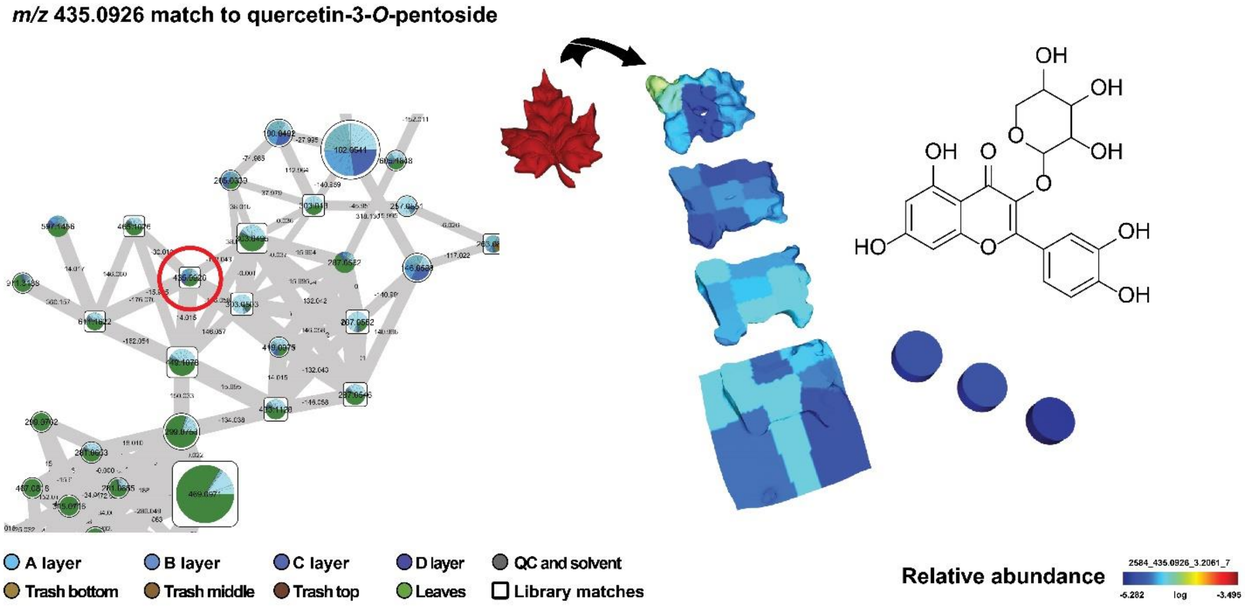


**Supplementary Fig. S6|Molecular cartography of quercetin-3-*O*-pentoside detected in *A. texana* fungus garden.** In this figure, the detected feature corresponding to quercetin-3-*O*-pentoside (*m/z* 435.0926) is highly abundant in the plant material, while its aglycone (**Supplementary Fig. S7**), quercetin (m/z 303.0503) is highly abundant in the plant material as well as in the top layer of the fungal garden. Connected nodes correspond to structurally related molecules based on their spectral similarity. The annotation of quercetin-3-*O*-pentoside from GNPS libraries (cosine 0.96) corresponds to a level 3, as the stereochemistry of the sugar moiety cannot be determined by MS/MS data, according to the 2007 metabolomics standards initiative.[7]

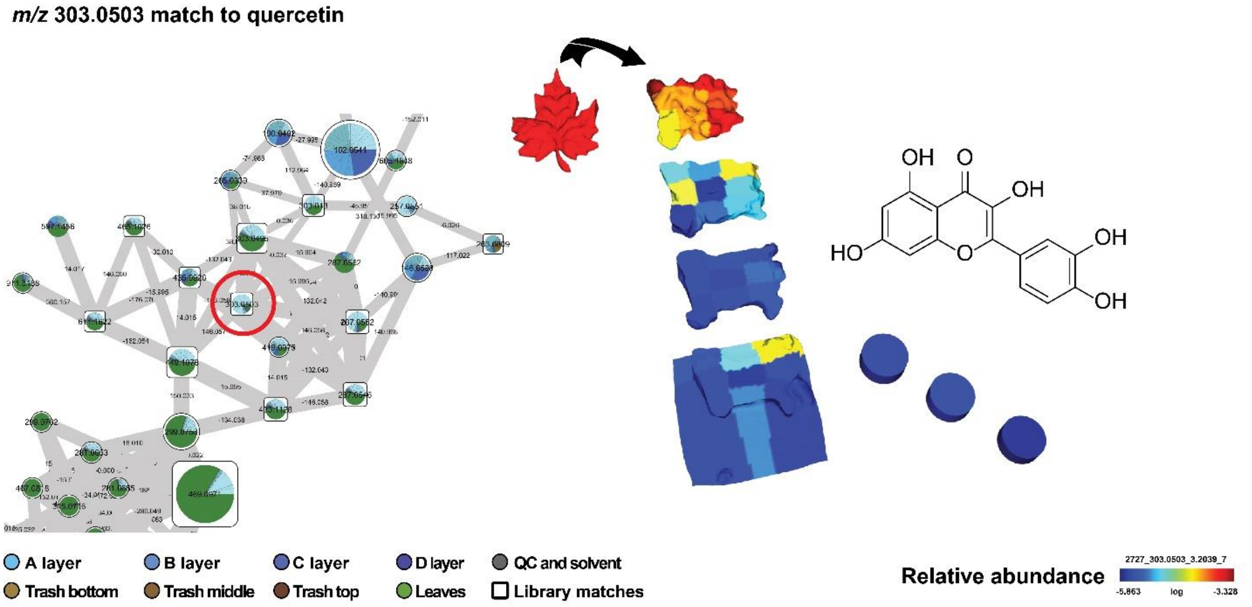


**Supplementary Fig. S7|Molecular cartography of quercetin detected in *A. texana* fungus garden.** In this figure, the detected feature corresponding to quercetin (*m/z* 303.0503) is highly abundant in the plant material as well as in the top layer of the fungus garden. Connected nodes correspond to structurally related molecules based on their spectral similarity. The annotation of quercetin from GNPS libraries (cosine 0.97) was confirmed by using a reference standard (**Supplementary Table S1**), then it corresponds to a level 1 match according to the 2007 metabolomics initiative.[7]

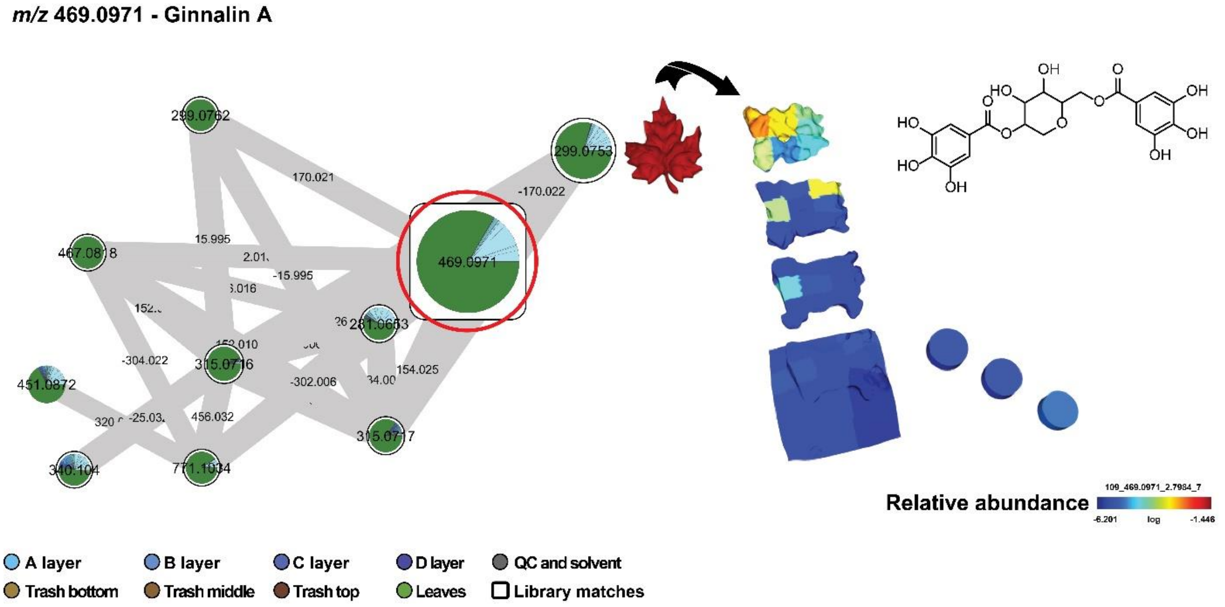


**Supplementary Fig. S8|Molecular cartography of ginnalin A detected in *A. texana* fungus garden.** In this figure, the detected feature corresponding to ginnalin A (*m/z* 469.0971) is highly abundant in the plant material as well as in the top layer of the fungus garden. The identification of ginnalin A based on spectral similarity to GNPS libraries (cosine 0.95) corresponds to annotation at level two according to the 2007 metabolomics standards initiative.[7]

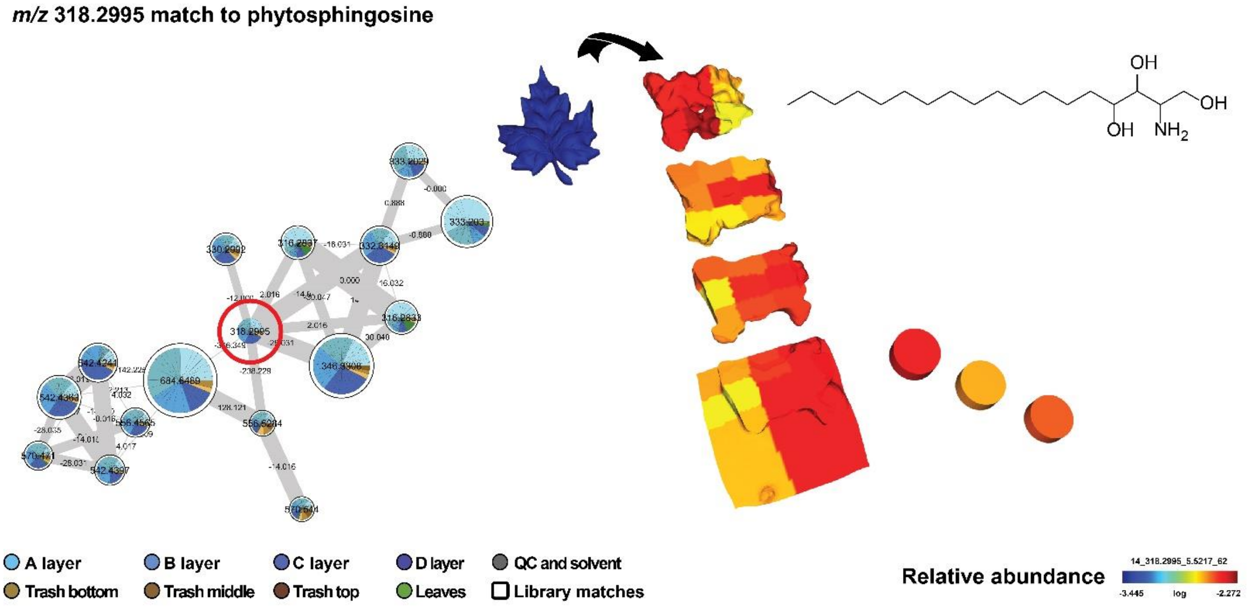


**Supplementary Fig. S9|Molecular cartography of phytosphingosine detected in *A. texana* fungus garden.** On the left: cluster of phytosphingosine-related molecules. On the right: spatial distribution of phytosphingosine (e.g. *m/z* 318.2995 $[M+H]^+$ match to phytosphingosine). Sphingosine-related metabolites have been involved in behavioural manipulation of ants by fungal pathogens.[8] The annotation of phytosphingosine from GNPS libraries (cosine 0.72) was confirmed by using a reference standard (**Supplementary Table S1**), corresponding to a level 1 match according to the 2007 metabolomics initiative.[7]

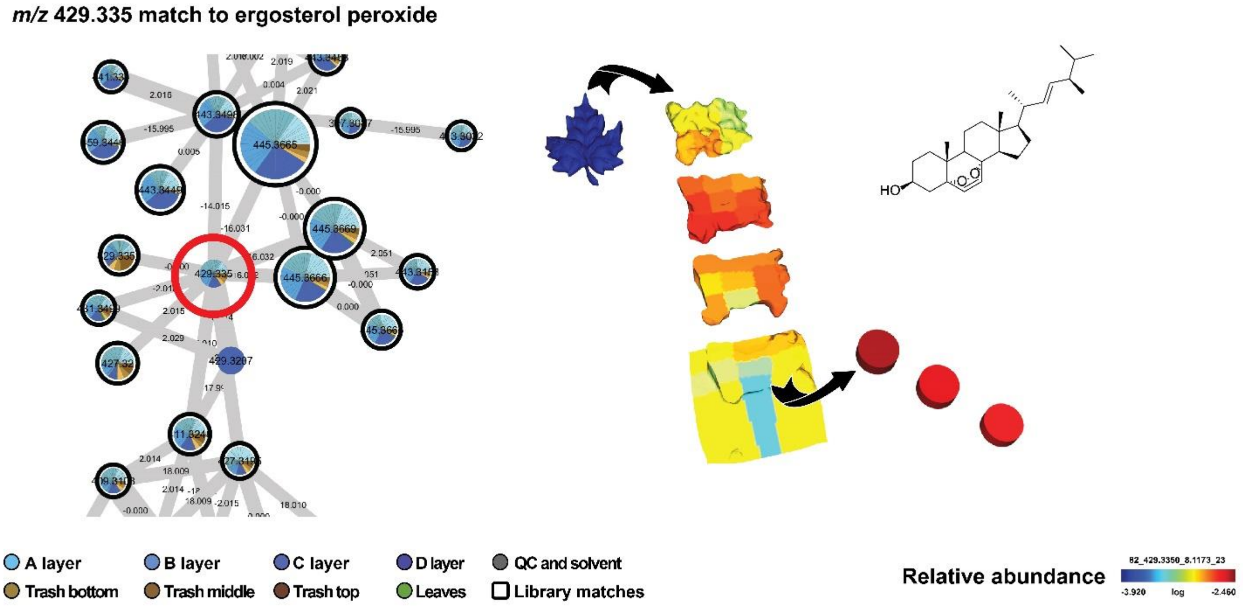


**Supplementary Fig. S10|Molecular cartography of ergosterol peroxide detected in *A. texana* fungus garden.** On the left: zoomed-in cluster of sterol-related molecules. On the right: spatial distribution of ergosterol peroxide (*m/z* 429.335 $[M+H]^+$). Although sterols are widely distributed in nature, and also present in fungi,[9] the relative high abundance in the trash material suggest they might be involved in chemical labeling of discarded material or might be a sub product of fungal decomposition potentially mediated by the microbiota associated with the waste material, as it has been reported previously.[10] The annotation of ergosterol peroxide from GNPS libraries (cosine 0.71) was confirmed by using a reference standard (**Supplementary Table S1**), a level 1 match according to the 2007 metabolomics initiative.[7]

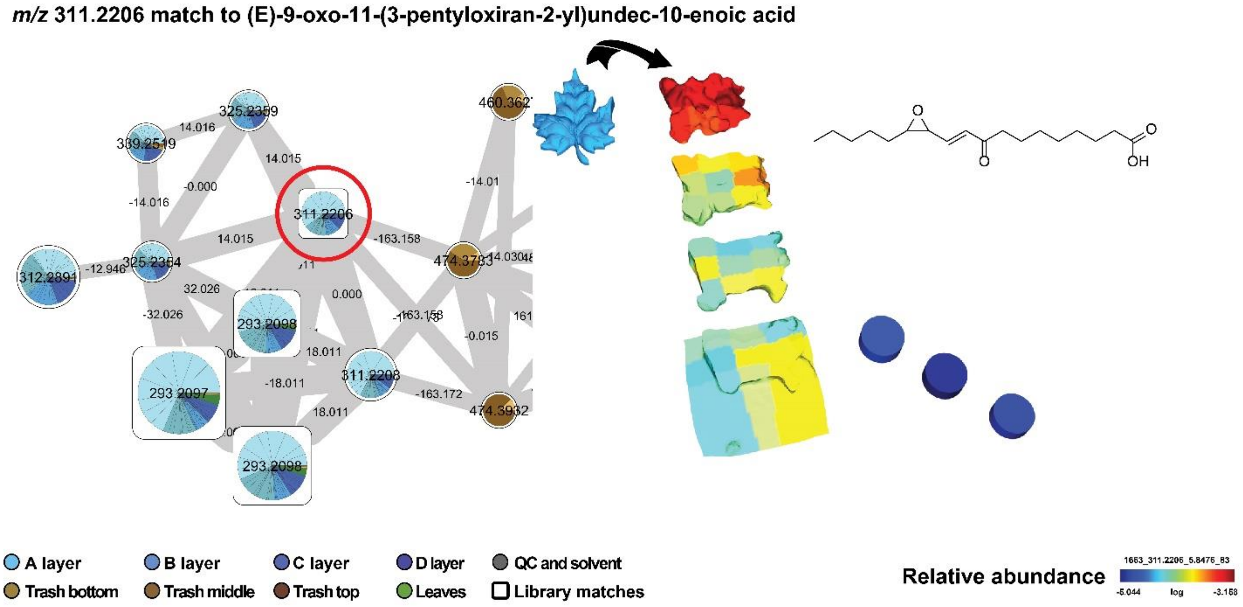


**Supplementary Fig. S11|Molecular cartography of (*E*)-9-oxo-11-(3-pentyloxiran-2-yl)undec-10-enoic acid detected in *A. texana* fungus garden.** On the left: zoomed-in cluster of oxylipin-related molecules. In the right: spatial distribution of oxylipin member of this cluster (e.g. *m/z* 311.221 $[M+H]^+$ match to (*E*)-9-oxo-11-(3-pentyloxiran-2-yl)undec-10-enoic acid, also known as trans-EKODE-(E)-Ib). Oxylipins are signaling molecules originated from the oxidation of polyunsaturated fatty acids and also involved in plant defense.[11,12] Common plant metabolites, these highly reactive molecules were detected in the fungus garden and trash material. Their presence is consistent with oxidation processes occurring in the fungal garden providing carbon sources available to the fungi but also they might play a role as signaling molecules for the present microbiota. The annotation of (*E*)-9-oxo-11-(3-pentyloxiran-2-yl)undec-10-enoic acid from GNPS libraries (cosine 0.86) was confirmed by using a reference standard (**Supplementary Table S1**), a level 1 match according to the 2007 metabolomics initiative.[7]

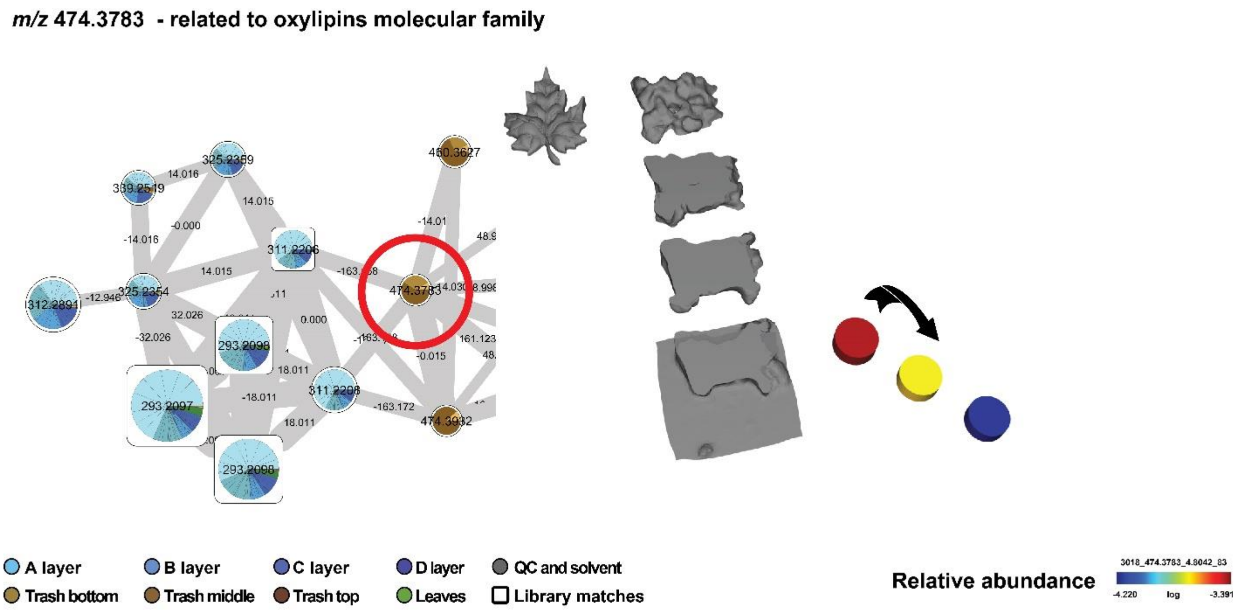


**Supplementary Fig. S12|Molecular cartography of putative oxylipin derivative detected in *A. texana* fungus garden.** On the left: cluster of oxylipin-related molecules. On the right: spatial distribution of an unknown oxylipin member of this cluster (e.g. *m/z* 474.3783). The arrow indicates that higher abundance of this feature was observed at the top of the waste material, and considerably decreasing to low abundance at the bottom of the waste material. It is possible the abundance is associated with accumulated biomass, higher at the top, although this needs further investigation. As mentioned in the previous figure (**Supplementary Fig. S11**), oxylipins are signaling molecules originated from the oxidation of polyunsaturated fatty acids and also involved in plant defense.[11,12] Common plant metabolites, these highly reactive molecules were detected in the fungus garden and trash material. Their presence is consistent with oxidation processes occurring in the fungus garden providing carbon sources available to the fungi but also they might play a role as signaling molecules for the present microbiota. This annotation, at the molecular family level, corresponds to a level 3 annotation according to the 2007 metabolomics standards initiative.[7]

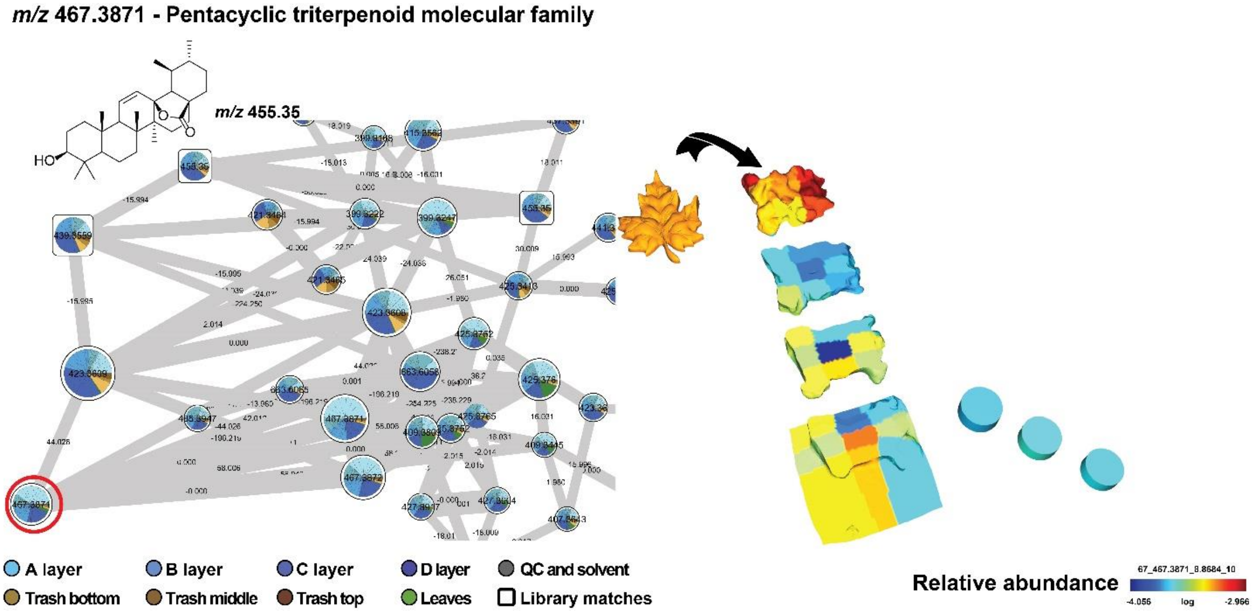


**Supplementary Fig. S13|Molecular cartography of triterpenoid derivative detected in** ***A. texana*** **fungus garden.** On the left: zoomed-in cluster of pentacyclic triterpenoid-related molecules. The chemical structure of 11,12-Dehydroursolic acid lactone (GNPS library match, cosine 0.79) corresponding to $C_{30}H_{46}O_3$ (*m/z* 455.35, $[M+H]^+$) is provided to illustrate this pentacyclic triterpenoid molecular family. On the right: spatial distribution of a member of this cluster ( *m/z* 467.3871 $[M+H]^+$) consistent with the molecular formula $C_{32}H_{50}O_2$. Although the distribution of this feature belonging to triterpenoid molecular family seems vague, it is evident its high abundance from the plant material and at the top layer of the fungus garden. This distribution contrasts with the distribution of the feature *m/z* 423.3609 shown in the next figure (**Supplementary Fig. S14**) and suggest this chemical transformation corresponding to the mass shift of 44.026 Da might occur in the fungal garden accumulating the product (feature *m/z* 423.3609) with high abundance at the bottom layers from the fungus garden and waste material. The annotation of the molecules shown at the molecular family level corresponds to a level 3 according to the 2007 metabolomics standards initiative.[7] The MS/MS of 11,12-Dehydroursolic acid lactone was found in 37 GNPS datasets comprising food, plants, lichen, mice, including human feces sample types.

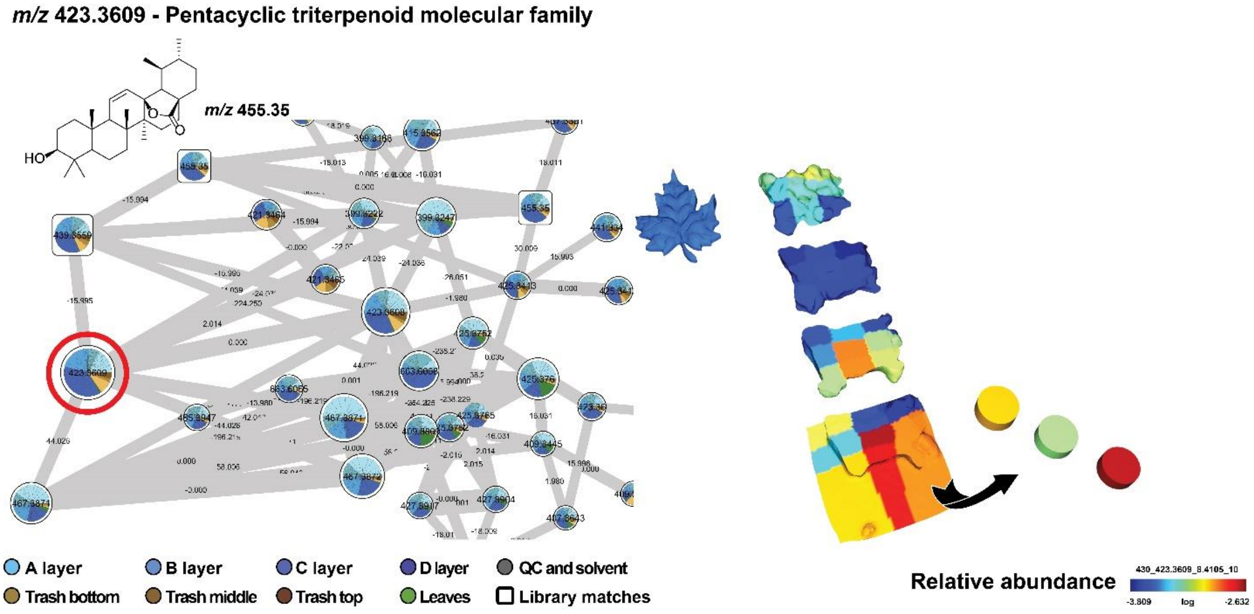


**Supplementary Fig. S14|Molecular cartography of triterpenoid derivative detected in *A. texana* fungus garden.** On the left: zoomed-in cluster of pentacyclic triterpenoid-related molecules. The chemical structure of 11,12-Dehydroursolic acid lactone (GNPS library match, cosine 0.79) corresponding to $C_{30}H_{46}O_3$ (*m/z* 455.35, $[M+H]^+$) is provided to illustrate this pentacyclic triterpenoid molecular family. On the right: spatial distribution of a member of this cluster (*m/z* 423.3609 $[M+H]^+$) consistent with the molecular formula $C_{30}H_{46}O$. This figure shows the distribution of this molecular feature with high abundance at the bottom layers from the fungus garden and waste material. Additionally, the abundances are higher in the waste material when compared with the fungus garden. Similarly, as observed for steroid and oxylipins, it is possible that metabolic transformations of terpenoids might be mediated by microorganisms present in the waste material. The annotation of the molecules shown at the molecular family level corresponds to a level 3 according to the 2007 metabolomics standards initiative.[7]

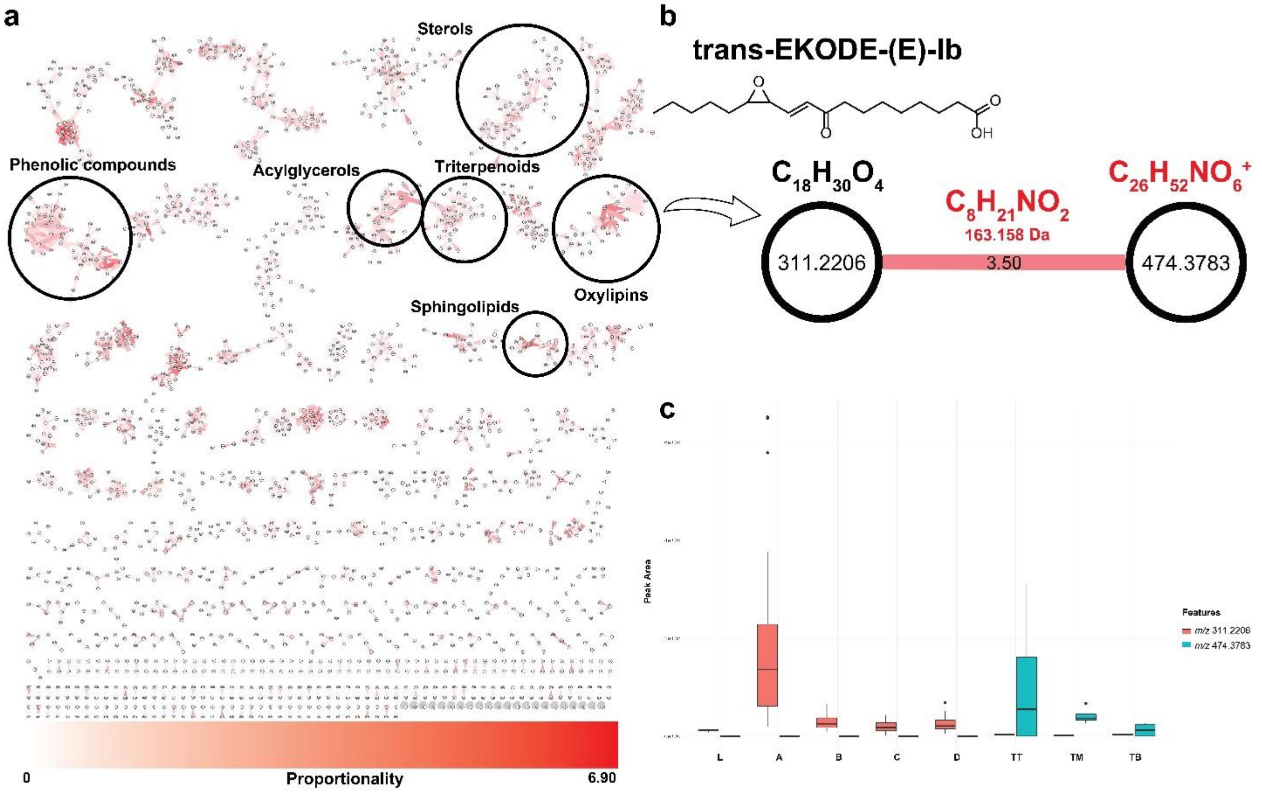


**Supplementary Fig. S15|Chemical proportionality of oxylipins molecular family calculated between the bottom layer of the fungus garden and top layer of the trash material. a.** Chemical features are highlighted from a molecular network based on their high proportionality score; **b.** The network pair *m/z* 311.2206 and *m/z* 474.3783 shows a mass shift of 163.158 Da putatively corresponding to $C_8H_{21}NO_2$. The annotation of the feature corresponding to *m/z* 311.2206 (*E*)-9-oxo-11-(3-pentyloxiran-2-yl)undec-10-enoic acid from GNPS libraries (cosine 0.86) was confirmed by using a reference standard (**Supplementary Table S1**), a level 1 match according to the 2007 metabolomics initiative.[7] The feature corresponding to node *m/z* 474.3783 (proposed molecular formula $C_{26}H_{52}NO_6^+$, 1.3 ppm error) was associated only to trash material while node *m/z* 311.2206 was detected in the bottom layer of the fungus garden and at low abundance in the trash material; **c.** The boxes represent the 25%, 50%, and 75% quartile and the whiskers extend ±1.5 times the interquartile range, and the jitters represent samples.

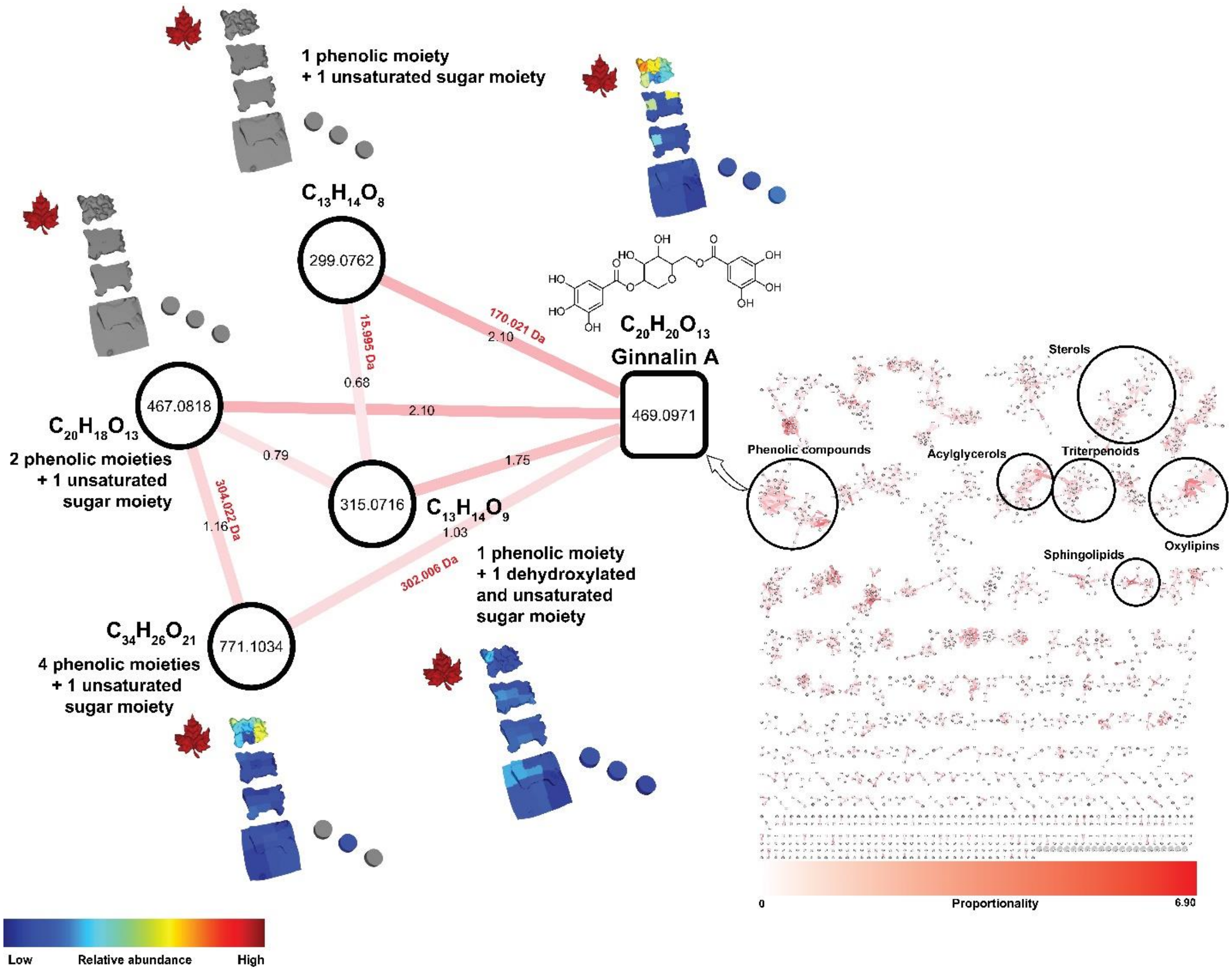


**Supplementary Fig. S16|Potential transformations of phenolic compounds related to ginnalin A.** Chemical features are highlighted from a molecular network based on their high proportionality score (red edge, edge label in black). The spatial distribution of the highlighted features by the proportionality approach are also shown in the 3D model of the *Atta texana* fungus garden (relative abundance shown as the color code in the figure). The feature corresponding to *m/z* 469.0971 annotated as ginnalin A (cosine 0.95), a bioactive phenolic compound,[13] was identified as a potential product of chemical transformation. Based on the information regarding the mass shifts (red labels, in Daltons) of the connected nodes and the similarity of fragmentation spectra (MS/MS), structural modifications were suggested. All the fragmentation spectra from the connected nodes in the molecular network shown in the figure shared the *m/z* 153.02 base peak corresponding to the phenolic moiety (gallic acid) indicating that a putative double bond is located in the sugar moiety. Ginnalin A was detected in the leaves as well as in the fungus garden while the features of *m/z* 299.0762 and *m/z* 467.0818 were only detected in the plant material. This suggest that ginnalin A and related features, *m/z* 315.0716(7) and *m/z* 771.1034 might be produced in a higher proportion than *m/z* 467.0818 and *m/z* 299.0762 that reach undetectable levels in the fungus garden. The identification of ginnalin A based on spectral similarity to GNPS libraries (cosine 0.95) corresponds to annotation at level 2 according to the 2007 metabolomics standards initiative,[7] while the match of the related molecules shown are at the molecular family level, a level 3 annotation.[7]

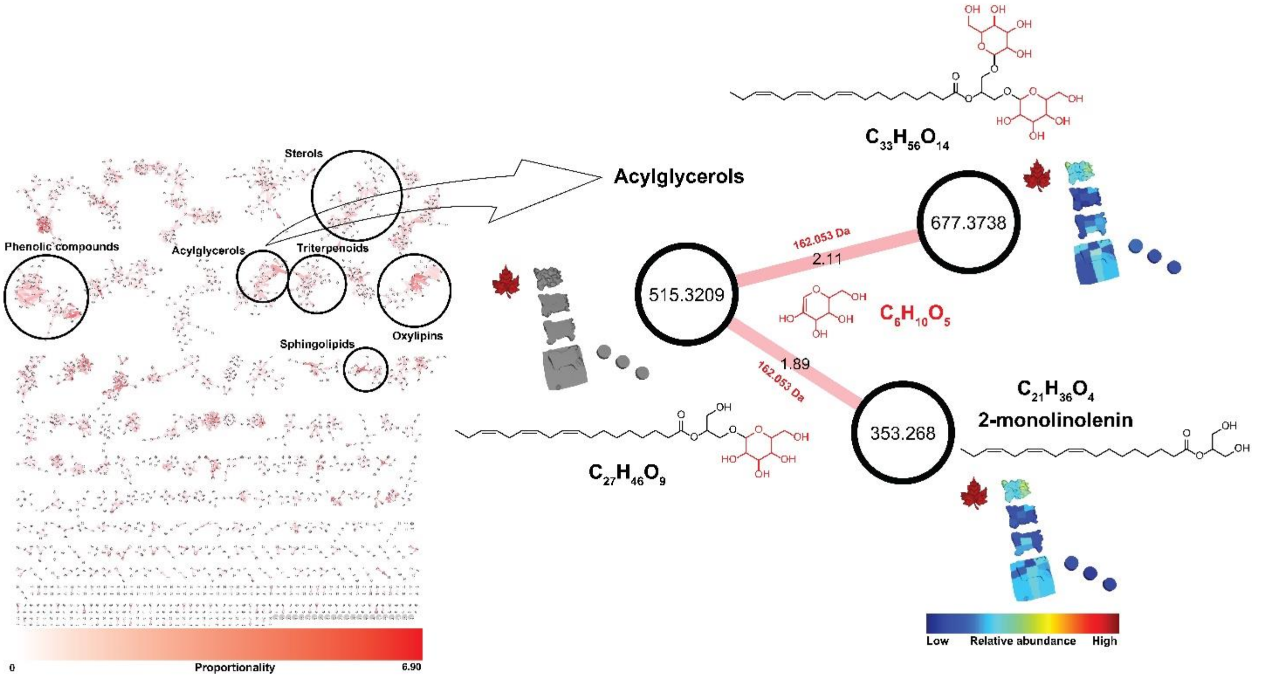


**Supplementary Fig. S17|Putative chemical modifications on acylglycerol derivatives highlighted by proportionality in *Atta texana* fungus garden.** A molecular family of acylglycerol compounds (left) are highlighted from a molecular network based on their high proportionality score. The feature corresponding to *m/z* 515.3209 was associated only to plant material while the feature *m/z* 677.3751 was detected across the samples, including fungus garden and trash material. The mass difference from this network pair was 162.053 Da corresponding to a molecular formula $C_6H_{10}O_5$, which is consistent with a sugar moiety. This example indicates that these compounds are involved in chemical transformation (gain of 162.053 Da) that occurs mainly in the fungus garden. For the structures shown, it was not possible to isolate sufficient quantities to analyze by NMR and therefore MS/MS matches to reference spectra was the only strategy to gain insight into the structural modifications. The identification of monolinolenin based on spectral similarity to GNPS libraries (cosine 0.75) corresponds to annotation at level two according to the 2007 metabolomics standards initiative,[7] while the match of the related molecules shown are at the molecular family level, a level three annotation.[7] For the monolinolenin derivatives, although chain length and number of double bonds are known, the location of the double bonds, cis or trans configuration of the double bonds or stereochemistry of the saccharide are not.

### Online methods

#### General overview of workflows applied in this study

To gain further information regarding chemical transformations occurring in leaf-cutter ant fungus gardens we deconstructed a laboratory maintained *Atta texana* fungus garden **(Fig. 1, Supplementary Fig. S1**). We applied spectral alignment using molecular networking,[14,15] spatial distribution of molecular features using molecular cartography,[16] and annotation of MS/MS spectra via GNPS workflows, including MolNetEnhancer,[1,3,15] to identify molecular signatures from the fungus garden samples. False discovery rate (FDR) for the compound annotations was estimated using the Passatutto decoy-based method.[17] We further introduce the concept of Chemical Proportionality (proportionality) using a modification to Meta-Mass Shift analysis recently introduced,[18] by considering also the abundances of the detected molecules, to quickly highlight features (metabolites) potentially involved in chemical transformations. Samples were prepared for untargeted profiling via reverse-phase liquid chromatography-tandem mass spectrometry (LC-MS/MS). Data were collected using data dependent acquisition mode and fragmenting the five most abundant precursor ions. Feature detection was performed using the open source software MZmine2.[19,20] Spectral alignments of the acquired MS/MS spectra and 3D visualization was performed using the Global Natural Products Social Molecular Networking (GNPS) platform.[14–16] By combining these approaches, we obtained molecular signatures from the fungus garden that enabled us to reveal chemical transformations occurring in specific locations of this system. Capturing these chemical transformations from the continuous process of plant degradation to waste accumulation is a fundamental step to understand the influence of the chemistry on microbial communities inhabiting these systems.[10]

### *Atta texana* fungus garden

The colony was maintained under laboratory conditions following standard methods for maintaining living colonies in the laboratory,[21] at the department of Molecular & Cell Biology at the University of Connecticut, laboratory of Prof. Jonathan Klassen. Briefly, colony JKH000189 was collected from Clear Creek Wildlife Management Area, LA (31.04990, -93.40245) under Louisiana Department of Wildlife and Fisheries Permit WL-Research-2016-10. Colony JKH000189 was maintained in plastic chambers where ants have access to fresh maple leaves (primarily *Acer platanoides*, but also *A. rubrum* and *A. saccharum*) provided every three days, as well as to an empty chamber that ants use as a waste container (**Supplementary Fig. S1**). Deconstruction of fungus garden, sample preparation, as well as data analysis were performed as follows:

### *Atta texana* fungus garden deconstruction

Three times, a section of the same laboratory maintained fungal garden was analyzed by this representative protocol. A 10x10x10 cm of fungus garden was removed from the chamber and scanned for creation of a virtual 3D image using Structure Sensor Mark II and Structure app [https://structure.io/structure-sensor/mark-ii]. The virtual 3D model was created using Meshmixer [http://www.meshmixer.com] software and the coordinates for *´ili’* visualization were obtained using Meshlab software [https://www.meshlab.net/]. Then, the piece of fungus garden was spliced in four layers and each layer was further divided into nine pieces of approximately 3x3x3 cm (or 27 $cm^3$). Each ~27$cm^3$ sample was homogenized by vortexing. Approximately 100 mg of each sample, including plant material (maple leaves) and waste material were extracted three times with 2:1 DCM:MeOH, sonicated for 10 minutes, dried under a stream of gaseous nitrogen and shipped to the Dorrestein Laboratory at UC San Diego for LC-MS/MS acquisition. The three data sets showed similar results. The molecular networking analysis can be accessed through the following links: first deconstruction, second deconstruction and third deconstruction. A short video showing the spatial distribution of the detected molecules from the third deconstruction of the *Atta texana* fungus garden can be accessed through the following link: **Supplementary Movie S1 [https://youtu.be/J-Ma_RNj6qs]**. Briefly, using the feature table of feature abundances from LC-MS/MS data (See **Feature-based molecular networking** described below) and the virtual 3D model we show the distribution of each signature detected in the fungus garden using the *´ili'* visualization (https://ili.embl.de/).

**Annotation of detected features**

Automatic annotation of detected molecules, via GNPS workflows,[1,3,15] were manually confirmed. Representative compounds from the main chemical classes discussed in the main text were confirmed by using reference standards, level 1 match according to the 2007 metabolomics initiative.[7] The following standards were used to confirm annotations at level 1 using spectral match, accurate mass and retention time: (10E)-9-Oxo-11-(3-pentyl-2-oxiranyl)-10-undecenoic acid (Cayman Chemicals Company, Inc.); quercetin (VWR International, LLC); quercitrin (Fisher Scientific), ergosterol peroxide (Carbosynth LLC), kaempferol (VWR International, LLC) and phytosphingosine (Sigma-Aldrich). See **Supplementary Table S1**:

**Supplementary Table S1.** Reference standards used to confirm annotated members of molecular families detected from *Atta texana* fungus garden.

| Reference compound | Detected *m/z* $[M+H]^+$ | Calculated *m/z* $[M+H]^+$ | Error (ppm) | Molecular Formula | Accurate Mass | CAS | Retention time (min) |
|---|---|---|---|---|---|---|---|
| Phytosphingosine | 318.2995 | 318.3003 | 2.4 | $C_{18}H_{39}NO_3$ | 317.2924 | 554-62-1 | 5.52 |
| EKODE (E)-9-oxo-11-(3-pentyloxiran-2-yl)undec-10-enoic acid | 311.2212 | 311.2217 | 1.6 | $C_{18}H_{30}O_4$ | 310.2139 | 478931-82-7 | 5.83 |
| Quercetin | 303.0498 | 303.0499 | 0.4 | $C_{15}H_{10}O_7$ | 302.0421 | 117-39-5 | 3.35 |
| Kaempferol | 287.0545 | 287.055 | 1.8 | $C_{15}H_{10}O_6$ | 286.0472 | 520-18-3 | 3.55 |

| | | | | | | | |
|---|---|---|---|---|---|---|---|
| Ergosterol peroxide | 429.3355 | 429.3363 | 1.9 | $C_{28}H_{44}O_3$ | 428.3285 | 2061-64-5 | 8.13 |
| Quercitrin | 449.1071 | 449.1078 | 1.6 | $C_{21}H_{20}O_{11}$ | 448.1000 | 522-12-3 | 3.36 |

**LC-MS/MS conditions**

Samples were resuspended in 100% methanol containing 2µM sulfamethazine as internal standard and LC-MS/MS analysis was performed in an UltiMate 3000 UPLC system (Thermo Scientific) using a Kinetex 1.7 ųm C18 reversed phase UHPLC column (50 X 2.1 mm) and Maxis Q-TOF mass spectrometer (Bruker Daltonics) equipped with ESI source. The column was equilibrated with 5% solvent B (LC-MS grade acetonitrile, 0.1% formic acid) for 1 min, followed by a linear gradient from 5% B to 100% B in 8 min, held at 100% B for 2 min. Then, 100%–5% B in 0.5 min and maintained at 5% B for 2.5 min at a flow rate of 0.5 mL/min throughout the run. MS spectra were acquired in positive ion mode in the range of 100-2000 m/z. A mixture of 10 mg/mL of each sulfamethazine, sulfamethizole, sulfachloropyridazine, sulfadimethoxine, amitriptyline, and coumarin was run after every 96 injections for quality control. An external calibration with ESI-Low Concentration Tuning Mix (*m/z* 118.086255; 322.048121; 622.028960; 922.009798; 1221.990637; 1521.971475; 1821.952313) (Agilent technologies) was performed prior to data collection. An internal calibrant Hexakis(1H,1H,2H-perfluoroethoxy)phosphazene (CAS 186817-57-2) was used throughout the runs. The capillary voltage of 4500 V, nebulizer gas pressure (nitrogen) of 2 bar, ion source temperature of 200 $^{o}$C, dry gas flow of 9 L/min source temperature, spectral rate of 3 Hz for MS1 and 10 Hz for MS2 was used. For acquiring MS/MS fragmentation, the 5 most intense ions per MS1 were selected, MS/MS active exclusion parameter was enabled, set to 2 and to release after 30 s, precursor ion was reconsidered for MS/MS if current intensity/previous intensity ratio >2. Advanced stepping function was used to fragment ions according to the **Supplementary Table S2** settings.

**Supplementary Table S2.** Instrument settings for data-dependent acquisition of *Atta texana* fungus garden samples.

| Time | Collision RF | Transfer Time | Collision |
|---|---|---|---|
| 0 | 450.0 | 70.0 | 125 |
| 25 | 550.0 | 75.0 | 100 |
| 50 | 800.0 | 90.0 | 100 |
| 75 | 1100.0 | 95.0 | 75 |

CID energies for MS/MS data acquisition were used according to the following settings:

| Type | Mass | Width | Collision | Charge State |
|---|---|---|---|---|
| Base | 100.00 | 4.00 | 22.00 | 1 |
| Base | 100.00 | 4.00 | 18.00 | 2 |
| Base | 300.00 | 5.00 | 27.00 | 1 |
| Base | 300.00 | 5.00 | 22.00 | 2 |
| Base | 500.00 | 6.00 | 35.00 | 1 |
| Base | 500.00 | 6.00 | 30.00 | 2 |
| Base | 1000.00 | 8.00 | 45.00 | 1 |
| Base | 1000.00 | 8.00 | 35.00 | 2 |
| Base | 2000.00 | 10.00 | 50.00 | 1 |
| Base | 2000.00 | 10.00 | 50.00 | 2 |

The mass of internal calibrant was excluded from the MS/MS list using a mass range of *m/z* 621.5–623.0.

### Feature-based molecular networking

Feature finding was performed with the open source MZmine software[19] version 2.38 using the settings provided in **Supplementary Table S3**. These pre-processing steps generated the .mgf file and quantification table to be used in the GNPS feature-based molecular network.

**Supplementary Table S3.** Pre-processing settings for feature detection using MZmine2 of LC-MS/MS acquired data from *Atta texana* fungus garden samples.

| | | |
|---|---|---|
| Mass detection | MS1 | 1.0E4 |
| | MS2 | 1.0E2 |
| Chromatogram building | Min time spam | 0.01 min |
| | Min height | 3.0E4 |
| | Tolerance | 25 ppm |
| Deconvolution [Baseline cut-off algorithm] | Min Peak height | 1.0E4 |
| | Peak duration range | 0.01-1.0 min |
| | Baseline level | 1.0E2 |
| | m/z range for MS2 scan pairing | 0.01 Da |
| | RT range for MS2 scan pairing | 0.3 min |
| Isotopic peak grouper | m/z tolerance | 25 ppm |
| | RT tolerance | 0.2 min |
| | Max charge | 2 |
| Alignment [Join Aligner] | m/z tolerance | 25 ppm |
| | Weight for m/z | 75 |
| | Weight for RT | 25 |
| | RT tolerance | 0.2 min |
| | RT correction | Checked |
| Gap filling [Peak finder] | Intensity tolerance | 1% |
| | m/z tolerance | 25 ppm |
| | RT tolerance | 0.2 min |
| | RT correction | checked |
| Peak filter | Peak area | 1.0E4-1.0E7 |
| Peak Row Filtering to export .mgf file to GNPS | Minimum peaks in a row | 2 |
| | RT | 1.00-14.00 min |

**False discovery rate (FDR) - Passatutto**

Passatuto decoy-based method[17] was used to estimate FDR for the annotations with the library match settings used for spectral identification: FDR < 0.072 was obtained for the obtained annotations using a minimum matching peaks of 6. **GNPS job link [Passatutto]:** 20190815_Atta_3D_metadata_coordinates_FBMNv125_Passatutto_ST05-PMT2-IT05. FDR < 0.16 was obtained for the obtained annotations using a minimum matching peaks of 5. **GNPS job link [Passatutto]:** MSV000082636_Passatutto_MMP5_ST05-PMT2-IT05

**Feature-based molecular network for deconstruction of *A. texana* fungus garden. Description:** A molecular network was created with the feature based molecular networking workflow (https://ccms-ucsd.github.io/GNPSDocumentation/featurebasedmolecularnetworking/) on the GNPS website (http://gnps.ucsd.edu). The data was filtered by removing all MS/MS fragment ions within +/- 17 Da of the precursor *m/z*. MS/MS spectra were window filtered by choosing only the top 6 fragment ions in the +/- 50Da window throughout the spectrum. The precursor ion mass tolerance was set to 0.02 Da and a MS/MS fragment ion tolerance of 0.02 Da. A network was then created where edges were filtered to have a cosine score above 0.7 and more than 6 matched peaks. Further, edges between two nodes were kept in the network if and only if each of the nodes appeared in each other's respective top 10 most similar nodes. Finally, the maximum size of a molecular family was set to 100, and the lowest scoring edges were removed from molecular families until the molecular family size was below this threshold. The spectra in the network were then searched against GNPS' spectral libraries. The library spectra were filtered in the same manner as the input data. All matches kept between network spectra and library spectra were required to have a score above 0.7 and at least 6 matched peaks. **GNPS**

**GNPS FBMN job link:** MSV000082636_3DAtta_metadata_coordinates_FBMN

**In silico annotation of detected features from *A. texana* fungus garden using GNPS workflows.**

**Description:** Network Annotation Propagation (NAP)[3] was performed via the GNPS platform. The following parameters were used: 10 first candidates for consensus score, 15 ppm accuracy for exact mass candidate search, positive acquisition mode, 0.5 cosine value to subselect inside a cluster; fusion results were used for consensus, search only for $[M+H]^+$ adduct type; 10 maximum candidate structures in the graph. The following databases were included in the search: Dictionary of Natural Products (DNP), Super Natural II (SUPNAT),[22] GNPS and Chemical Entities of Biological Interest (ChEBI).[23] GNPS NAP job link: **MSV000082636_3DAtta_metadata_coordinates_FBMN_NAP** Automatic workflows for peptide analogues was performed using VarQuest[24] in GNPS and can be accessed using the following link: **MSV000082636_3DAtta_metadata_coordinates_FBMN_VarQuest.** A complementary analysis for substructures present in the dataset was performed using the mass-to-motif MS2LDA[25] workflow in the

GNPS and can be accessed through the following link: MSV000082636_3DAtta_metadata_coordinates_FBMN_MS2LDA_MotifDB

**MolNetEnhancer for molecular network of deconstructed *A. texana* fungus garden. Description:** Molecular network annotations were increased via the MolNetEnhancer workflow[1] merged in the GNPS platform. The workflow merged *in silico* annotations from Network Annotation Propagation (NAP),[3] VarQuest[24] and MS2LDA[25] to provide structures annotations at the class level. **MolNetEnhancer job link:** https://gnps.ucsd.edu/ProteoSAFe/status.jsp?task=50dfaf589c8140bd85a8ca199db59de3

**MassIVE links**

The data sets used in this work were deposited in the online repository namely GNPS/MassIVE. The data set corresponding to molecular cartography of *A. texana* fungus garden [**MassIVE:** MSV000082636] can be accessed via the following link: https://gnps.ucsd.edu/ProteoSAFe/result.jsp?task=df2eb5792c84460b9413fa22af2d0d89&view=advanced_view

**Proportionality score**

The proportionality score calculated between two directly connected nodes across the entire molecular network was obtained using the following equation:

$$Proportionality = log\frac{N_{s1}/M_{s1}}{N_{s2}/M_{s2}}$$

Where $N_{S1}$ and $M_{S1}$ correspond to the peak area of the detected features N and M in sample S1, while $N_{S2}$ and $N_{S2}$ correspond to the peak area of the detected features N and M in sample S2. Samples S1 and S2 correspond to leaves (L), layers of fungus garden (A, B, C, D) and layers of trash (TT, TM and TB). A constant ($k$ = 1.0 $e^{-10}$) is added to each value to avoid any zero values to be included during the calculation. **Supplementary Table S8** (Chemical proportionality table calculated between each sample type: L:A, A:B; B:C; C:D; D:TT; TT:TM; TM:TB) and **Supplementary Table S9** (Chemical proportionality table calculated between Leaves vs entire dataset: L:A; L:B; L:C; L:D; L:TT; L:TM; L:TB) contain the calculated scores for the entire dataset reporting the maximum proportionality score and the compared samples (e.g., L:A meaning Leaves:Layer A according to the scheme in Figure 1b). Maximum proportionality scores are visualized in **Fig. 2-3** from the main text as well as **Supplementary Fig. S15-S17**.

**Statistical analyses.** Statistical analyses were performed using the MetaboAnalyst platform.[5,6] The peak intensity table obtained after preprocessing with MZmine2 software was uploaded to MetaboAnalyst [Samples in columns (unpaired)]. The uploaded data file contained 160 (samples) by 3717 features [peaks (m/z and RT) data matrix], eight groups were included in the table (leaves, layers of fungus garden (A, B, C and D) and trash (TT, TM and TB) material), 83 features with a constant or single value across samples were found and deleted. No missing values were found. Data filtering: interquartile range (IQR) was used to detect filter variables that were near-constant values. The data were normalized by using a reference feature. In this case the internal standard was used, sulfamethazine [$C_{12}H_{14}N_4O_2S$] corresponding to the *m/z* 279.0910, 2.55 min. Cube root transformation (cube root of data values) was performed to facilitate feature comparison (**Supplementary Fig. S3-S4**).

**Supplementary Tables S4 – S9** are provided as .csv files

**Supplementary Table S4.** PLSDA coefficient scores of deconstructed *Atta texana* fungus garden samples.

**Supplementary Table S5.** PLSDA loadings values of deconstructed *Atta texana* fungus garden samples.

**Supplementary Table S6.** PLSDA scores of deconstructed *Atta texana* fungus garden samples.

**Supplementary Table S7.** VIP scores for PLSDA model of deconstructed *Atta texana* fungus garden samples.

**Supplementary Table S8.** Chemical proportionality table [Leaves/Fungus garden/Trash layers]

**Supplementary Table S9.** Chemical proportionality table [Leaves vs entire dataset]
